# Several complementary methods of plant establishment are required for effective restoration of banksia woodland

**DOI:** 10.1101/2020.07.09.195131

**Authors:** Mark Brundrett, Anna Wisolith, Margaret Collins, Vanda Longman, Karen Clarke

## Abstract

New banksia woodland vegetation was established at two sites totalling 50 ha in the Perth region of Western Australia as part of an offset-funded project. Restoration methods included topsoil transfer (16 ha), planting of nursery-raised local provenance seedlings (46,000 seedlings over 39.5 ha) and direct seeding with machinery or by hand (16.5 ha) with treatments overlapped. Six years of rigorous monitoring revealed trends in plant diversity, density and cover and allowed comparison of vegetation structure and composition to reference sites. Of the 162 native plants recorded, 115 originated primarily from the topsoil seed bank and the remainder from planting and seeding. Native plant germination from topsoil peaked at 700,000 stems per ha in year 2, but there was very high attrition during extreme summer drought. By year 5, native perennials averaged 20,000 stems per ha, well above the target of 7,000, but there was high spatial variability in plant density with 1/3 of quadrats below target. Saplings of Banksia spp. (the dominant local trees) plateaued at 150-220 stems per ha due to high summer mortality. Native plant cover reached 20% and perennial weed cover stabilised at under 5% within 5 years. Several complimentary methods were required for successful restoration, since transferred topsoil established most of the plant diversity. However, trees required planting or seeding, due to their canopy stored seed. Direct seeding and planting without respread topsoil led to lower overall diversity and density, but higher tree density. Most completion criteria targets were reached after 5 years. Areas with respread topsoil are trending towards recovery as a banksia woodland, but areas with only planting and seeding are likely to remain a separate vegetation type. Evidence for resilience of restored areas was provided by abundant pollination and seed set and second-generation seedlings. We suggest it may be possible to restore banksia woodland despite major challenges due to unpredictable offset funding, climate, weeds, grazing, recalcitrant species, and seed availability, but long-term monitoring is required to confirm this.

## Introduction

Banksia woodlands of the Swan Coastal Plain in Western Australia are sclerophyllous plant communities with an overstory dominated by banksia species, especially *Banksia attenuata* and *B. menziesii*, along with eucalypts, especially *Eucalyptus todtiana* and *E. marginata* (Government of Western Australia 2000). The understory in these vegetation types has a high diversity of shrubs, herbs, geophytes, annuals, and sedges. Substantial areas of banksia woodland have been lost to urban development and the remnants face additional threats such as water table drawdown, *Phytophthora* dieback disease, weeds and increased fire frequency and intensity (Stenhouse et al. 2004, Fisher et al. 2006, Ramalho et al. 2014, Department of the Environment and Energy 2016). These threats have resulted in banksia woodlands on the Swan Coastal Plain being listed as a threatened plant community (Department of the Environment and Energy 2016). Thus, it is especially important to monitor population trends after habitat restoration for key species and identify processes which impact on the ecological sustainability of restored banksia woodland.

Australian governments may require organisations that clear substantial areas of banksia woodland to offset these biodiversity losses by funding restoration projects that aim to purchase, restore, or improve similar habitats elsewhere. Judging the effectiveness of these projects requires proponents to demonstrate that environmental benefits fully compensate for the losses of biodiversity which triggered the offset, but this information is often deficient (Gibbons & Lindenmayer 2007, Thorn et al. 2018). Thus, offset funded projects need to be accountable and auditable to counter criticism of their effectiveness (Maron & Louis 2018). In particular, judging the success of restoration projects requires outcomes to be reported relative to completion criteria which measure the composition, structure, and resilience of restored vegetation relative to reference ecosystems (McDonald et al. 2016).

The Banksia Woodland Restoration (BWR) project was established in 2011 as part of the Commonwealth’s ministerial conditions to offset the impacts of clearing 167 ha of banksia woodland at Jandakot Airport, Perth, Western Australia (Commonwealth of Australia 2010). The project was managed by the Department of Biodiversity, Conservation and Attractions (DBCA) to create new banksia woodlands and repair existing woodlands in the Perth Metropolitan Region. The BWR project activities reported here concern banksia woodland restoration with the following specific objectives: (a) comparing the efficiency and effectiveness of different restoration methods, (b) establishing a comprehensive monitoring program to measure outcomes relative to completion criteria, (c) use monitoring data to update species list for planting and seeding to maximise native plant diversity and cover (adaptive management), (d) develop solutions to factors limiting plant recruitment and survival and (e) provide advice to other similar projects. This paper describes outcomes of the first five years of this project relative to completion targets based on reference site data, as explained in the accompanying paper (Brundrett et al. submitted). We also provide data on overall vegetation cover trends using remote sensing data in the third paper of this series (van Dongen et al. submitted).

## Materials and Methods

The selection of restoration sites, reference site surveys and development of completion criteria are described in the first part of this study (Brundrett et al. submitted). As shown in Figure 1, the two main restoration areas were a 39-ha site at Anketell Road (AR) in Jandakot Regional Park and 11 ha at Forrestdale Lake (FL). The layout of the largest site (AR) is shown in Figure 1. The sites were initially open, weed-dominated fields with a long history of agricultural land use (Figure 2). Areas restored by topsoil transfer, planting and seeding from 2012 to 2016 at AR are shown in Figure 1. The locations of monitoring quadrats are also indicated.

**Figure 1.**
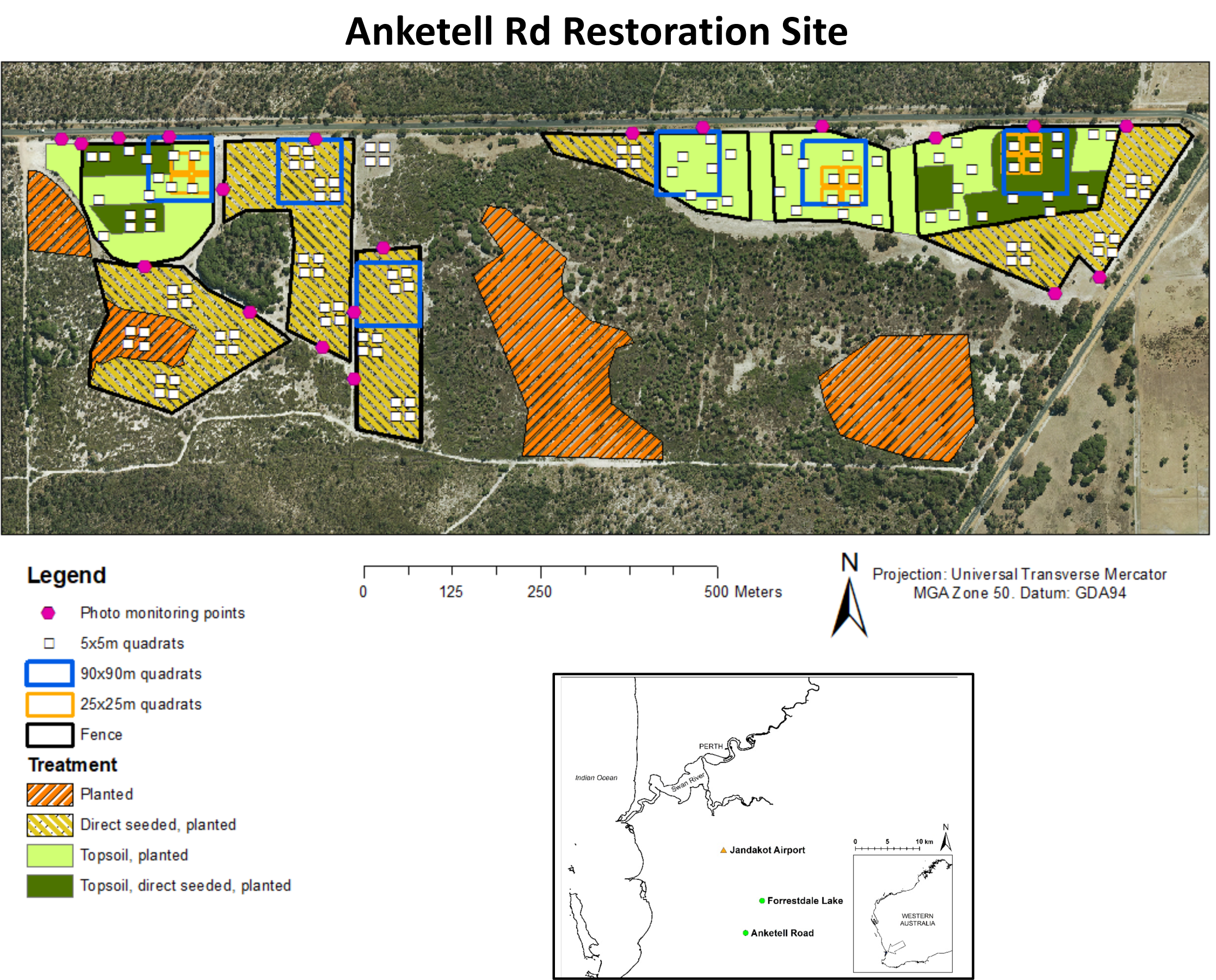
Aerial photograph of Anketell Road restoration site showing areas spread with topsoil from Jandakot Airport, fenced, direct seeded, and planted over the period 2012 to 2016. Monitoring points and quadrats are also shown.

**Figure 2.**
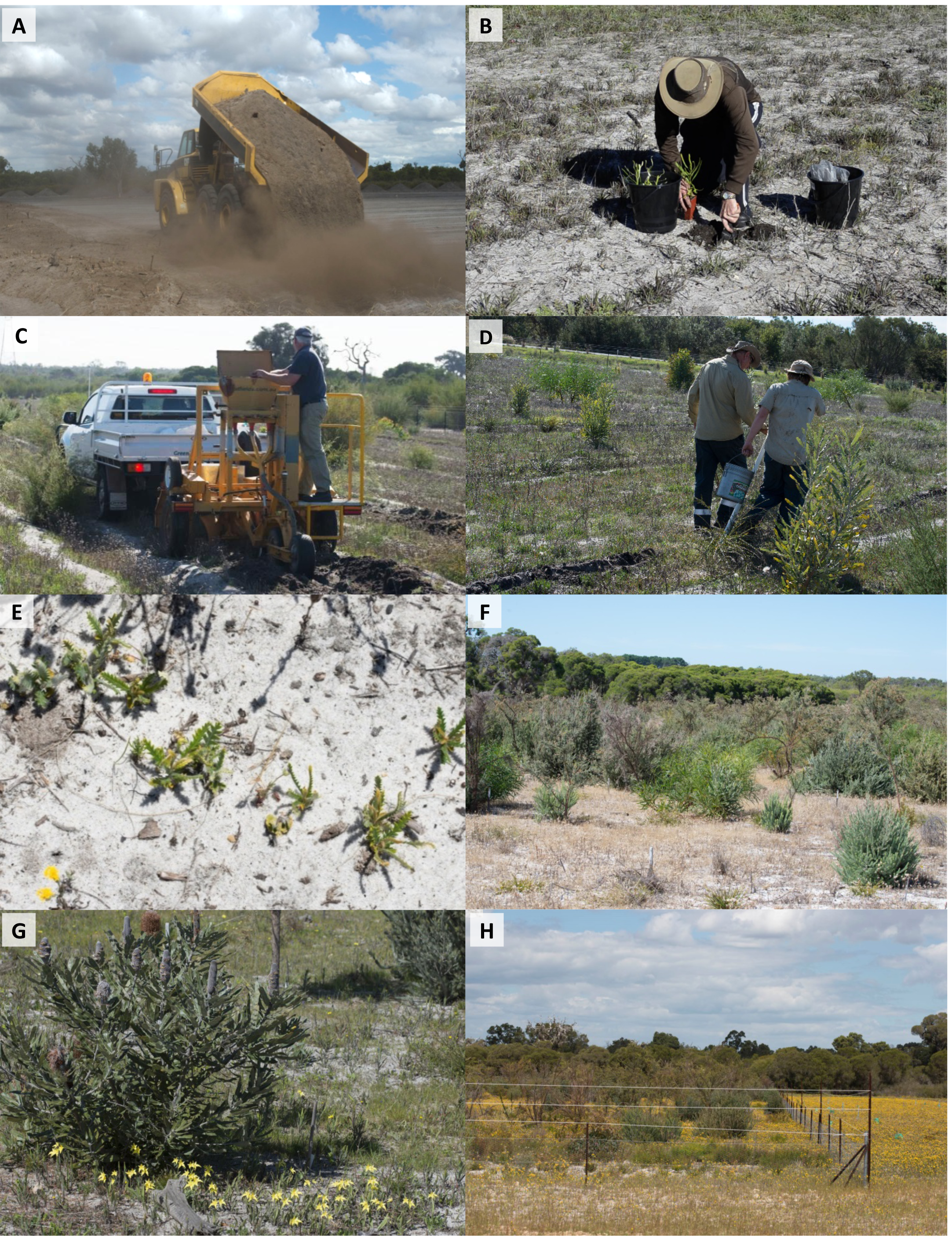
Restoration methods and outcomes at the Anketell Road restoration site. **A**. Spreading topsoil on open paddock in April 2012, after removal of a thin layer of the paddock’s weed infested topsoil **B**. Planting tubestock with DBCA staff and Aboriginal Trainees in 2015. **C**. Direct seeding using a seed drill at Anketell Road. **D**. Manual direct seeding in 2015. **E**. Seedlings resulting from the manual direct seeding, including 15 banksias and one Christmas tree (*Nuytsia floribunda*) in close proximity. **F**. Area with low survival of native plants in foreground and dense natives in background. **G**. Flowering of cowslip orchids (*Caladenia flava*) next to a 4-year old *Banksia menziesii* tree with seed. **H**. Very low survival of plants outside of the rabbit-proof fence (note white sight lines used to deter kangaroos).

### 1. Seed collections and nursery orders

Species considered unlikely to be present in the topsoil seed bank were the focus of seed collecting and nursery orders (see Table 2 in Brundrett et al. submitted). These included the majority of Carnaby’s cockatoo food plants and all trees which have canopy-stored seed. Local provenance seed collected for the BWR project included large quantities from areas about to be cleared at Jandakot Airport, as well as 50 other locations with similar soils and vegetation types. In total, 1,238 accessions of seed were collected from 164 species. Seed was stored in a refrigerated and humidity-controlled environment in DBCA’s Threatened Flora Seed Centre. Seed quality was assessed through germination testing, and field trials to test different germination and storage conditions.

**Table 1.** The size of areas that received topsoil, seed, or plants at the two restoration sites by year. Treatments overlapped in various combinations therefore total areas restored are not sums of each method.

| Restoration activity | Year | Anketell Rd (ha) | Forrestdale Lake (ha) | Total (ha) |
| --- | --- | --- | --- | --- |
| **Topsoil transfer** | **2012** | **11.5** | **4.5** | **16** |
| **Planting** | 2012 | 2.5 | 1.5 | 4 |
|  | 2013 | 9 | 7 | 16 |
|  | 2014 | 32 | 7.5 | 39.5 |
|  | 2015 | 26 | 3 | 29 |
|  | **TOTAL** | **32** | **7.5** | **39.5** |
| **Direct seeding** | 2012 | 2 | 0.5 | 2.5 |
|  | 2013 | 10 |  | 10 |
|  | 2014 | 12 |  | 12 |
|  | 2016 | 12.5 | 1 | 13.5 |
|  | **TOTAL** | **15.5** | **1** | **16.5** |
| **Fencing** | **TOTAL** | **23** | **3.5** | **26.5** |
| **Total site area** |  | **39** | **11** | **50** |

**Table 2.**
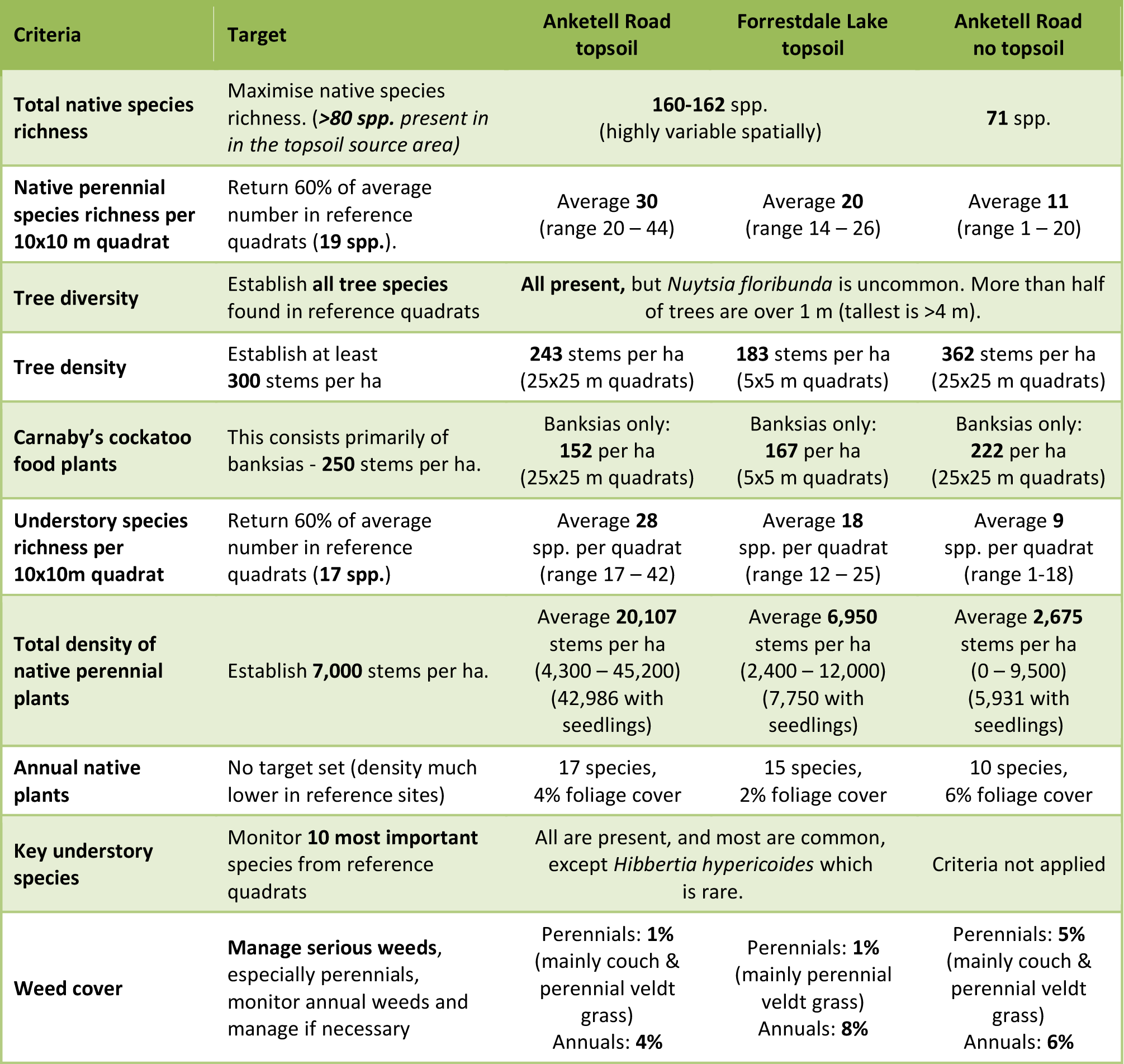
The status of restoration outcomes in spring 2017 relative to targets set based on the topsoil source reference site. These targets were not applied to areas at Anketell Road without topsoil but are shown for comparison.

**Table 3.** Potential advantages and disadvantages of different restoration methods in banksia woodland. Note that all methods also require effective grazing exclusion and weed management.

| Method | Advantages | Disadvantages |
| --- | --- | --- |
| **Recruitment from soil seed bank following topsoil transfer** | • High plant diversity and abundance is possible.<br>• Often includes species that cannot be easily grown in nurseries.<br>• May reduce weeds by covering existing soil. | • Some key species are likely to be missing.<br>• Soil may contain weeds and/or disease organisms.<br>• Soil seedbank can lose viability if transferred when wet or stockpiled.<br>• Soil transport is expensive. |
| **Planting of nursery tubestock** | • Can be cost effective for many species.<br>• Often effective for trees and other dominant plants.<br>• Required for species absent from topsoil seedbank, in Banksia woodland this includes most trees and large shrubs. | • Planting is slow and laborious.<br>• High summer mortality is common.<br>• Seedlings must overcome problems with root system growth in confined spaces.<br>• Seeds are difficult to obtain for some species. |
| **Direct seeding using machinery or by hand** | • Relatively inexpensive and rapid method for large areas.<br>• Often effective for trees and other dominant plants.<br>• Initial weed cover reduced in furrows.<br>• Infill hand seeding can be used to supplement initial establishment.<br>• Seed mixes can be adjusted for different plant communities. | • High mortality or poor germination may occur.<br>• Plants establish in discrete rows with gaps where weeds can grow.<br>• Final diversity is often be low due to seed availability and competition by more aggressive species. |

Approximately 6.4 kg of seed from 49 species was sent to 4 nurseries to produce over 46,000 tubestock plants that were planted in restoration areas (Table 1). A further 1000 plants of 7 species that could not be easily propagated from seed were produced by clonal division at a specialist nursery (Table S1). We also supplied seed to restoration projects run by community groups and local governments (Brundrett et al. 2018).

### 2. Restoration practices

Topsoil from Jandakot Airport was harvested to a depth of 50 mm, transported to restoration sites, and spread to a uniform depth of either 50 or 100 mm in April-May 2012 (Fig. 2A). A thin layer of existing topsoil (5-10 cm) was removed prior to topsoil spreading to reduce the weed soil seed bank. At AR, 11.5 ha of upland areas received transferred topsoil and 4.5 ha at FL. Control of grazing animals required rabbit-proof fencing with a total enclosed area of 26.5 ha (Fig. 1, Table 1). Fences were 1-1.5 m high with a buried skirt about 30-50 cm deep x 30 cm wide to exclude rabbits, as well as two or three white plastic “sight lines” suspended 30-60 cm above the fence to deter kangaroos (Fig. 2H).

Weed species were ranked according to their invasiveness and competitive ability and initially management focussed on perennial veldt grass (*Ehrharta calycina*) using a grass-selective herbicide (Fusilade®). Other invasive perennial weeds were removed by hand (all bulbs, *Euphorbia, Pelargonium, Carpobrotus, Lupinus*, etc.). Veldt grass, pigface (*Carpobrotus edulis)* and couch grass (*Cynodon dactylon*) were sprayed annually with appropriate herbicides from 2012 to 2017.

Planting and seeding occurred from 2012 to 2016 (Table 1), with separate species lists for upland and ecotone dampland areas. Tubestock of perennial native plants was planted by staff, volunteers from community groups and contactors (Fig. 2B) primarily within the fenced areas. Direct seeding was used to revegetate areas where topsoil transfer was not possible (Fig. 1). A total of 16.5 ha at AR and FL were direct seeded by Greening Australia WA using seed drill technology from 2012-2016. Seeds were mixed with a wetting agent, fertiliser, sand, and vermiculite (Fig. 2C). To infill smaller areas, we developed a hand direct seeding method using seed mixed with bulking agents. A shallow furrow was dug with a hoe and then the seed mix funnelled through a 1 m plastic pipe while walking down the rows (Fig. 2D). The seeds were then covered with a thin layer of soil using a rake or hoe. In total, more than 63 kg of seed from over 80 species was used for direct seeding by both methods, including 30 kg of the very large seeds of the cycad *Macrozamia fraseri*.

### 3. Monitoring

Monitoring of restoration areas for comparison with the completion criteria targets (Table 2) required the use of very small quadrats for seedlings and annuals, medium sized quadrats for perennial plants and large quadrats for trees (Wisolith et al. 2017). This was due to very large differences in plant density which normally occur for these categories (Kent 2012). From 2012-2016, cover and density of all species were measured within 1×1 m quadrats arranged along transects. From 2014-2017, 5×5 m quadrats were used to monitor density and foliage cover of perennial natives and weeds, as well as the cover of annuals. In total there were 144 of these quadrats, with 80 in AR and FL areas restored with topsoil, and 64 in AR areas with direct seeding and planting only, as shown in Figure 1. Data from groups of four quadrats were used to create virtual 10×10 m quadrats for comparison with reference sites.

We were able to identify both seedlings and adults of native plants using a photographic catalogue (Brundrett and Wisolith 2020) and reference herbarium. Unidentified seedlings were tagged to allow later confirmation of their identity. In some cases, collected specimens were identified using the resources of the Western Australian Reference Herbarium and by consultation with experts. Detailed images of all species present were also made available online (www.flickr.com/groups/banksia_plants).

Quadrats 25×25 m are used to measure tree density, flowering, and seed production (Fig. 1). The basal diameter and height of trees over 1 year old was also measured annually in both 5×5 and 25×25 m quadrats. Photo-monitoring points were established in all restoration areas at 20 locations and monitored biannually to illustrate changes in vegetation structure. We also took downward photos of 1×1 m quadrats to help measure plant cover (see Brundrett et al. 2018). Incremental changes in the overall cover of vegetation was also monitored by aerial photographs and remote sensing (van Dongren et al. submitted). Changes in floristic composition at revegetation sites over time relative to reference sites were investigated using multivariate analysis of plant cover data. Overall results for both the AR and FL sites are presented, but discussion of outcomes primarily concern the larger AR site.

## Results

### 1. Restoration outcomes

Plant survival and growth during the first five years of restoration at AR and FL were seriously impacted by periods of exceptionally severe drought in autumn, winter, and/or late spring in most years of the project (Fig. 3). This caused severe mortality of seedlings from topsoil and planted nursery-raised seedlings at the restoration sites in 2012, 2013 and 2015 (Fig. 3). Despite the high attrition of seedlings, foliage cover rapidly increased over time (Figs. 4, 5) but remained patchy after 5 years. By autumn 2017, there was twice as much cover at AR (21%) as at FL (12%), for reasons explained below. Weed cover stabilised at 6% at AR and 4% at FL (Fig. 4). Areas restored without topsoil at AR had a lower cover of natives (8%), because restoration work began two years later and consisted of direct seeding and planting only (Fig. 4). Native shrubs dominated native plant cover at both sites (17% in AR topsoil areas), trees were just over 2%, while herbs, sedges and grasses together were only 1% of cover. Most of the increase in perennial plant cover occurred over summer due to growth of shrubs and trees, while weeds were most active in winter. There was a complex gradient in species importance measured as cover, where a few large and common species were most dominant, but the majority of species present were much smaller or uncommon (Fig. 5).

**Figure 3.**
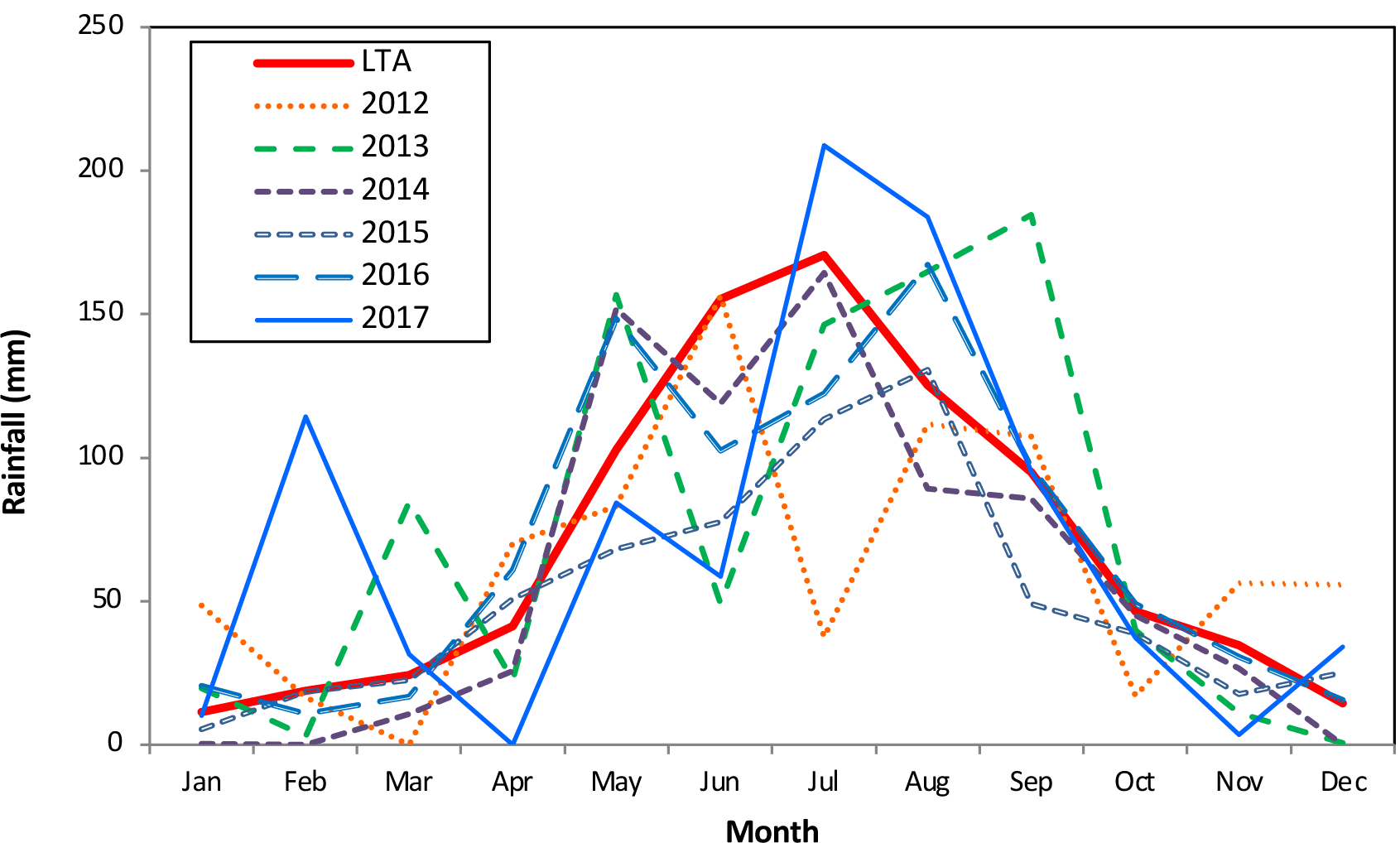
Monthly rainfall (mm) at the Anketell Road weather station over the period 2012 to 2017 (www.bom.gov.au). Red line indicates the long-term average (LTA) for 1985-2017. Mean annual rainfall for this period is 841 mm.

**Figure 4.**
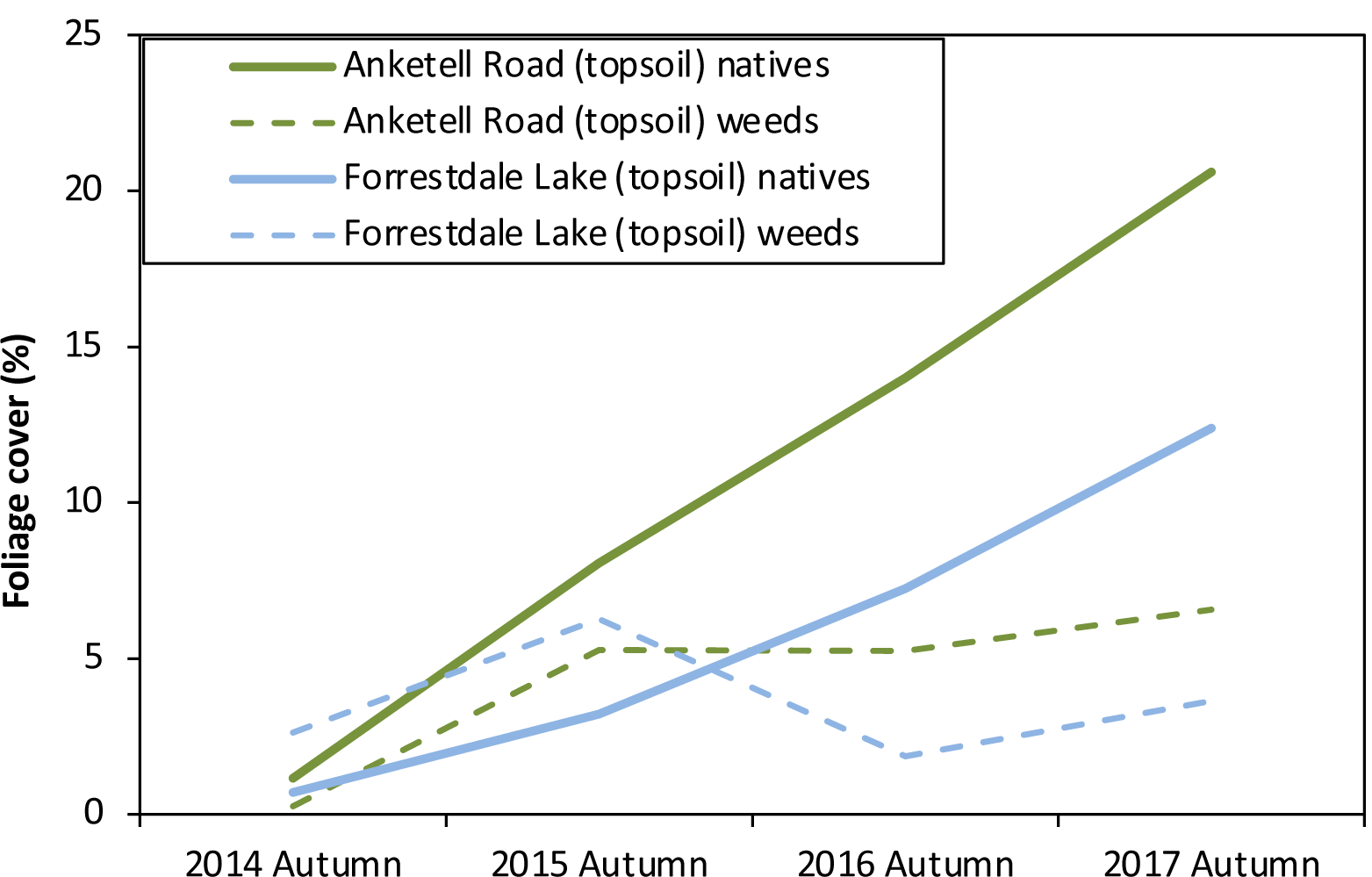
Changes in foliage cover for weeds and native plants at two restoration sites. Results show cover in the 5×5m quadrats over 2014-2017. Cover of natives steadily increases over time while weed cover remains relatively stable.

**Figure 5.**
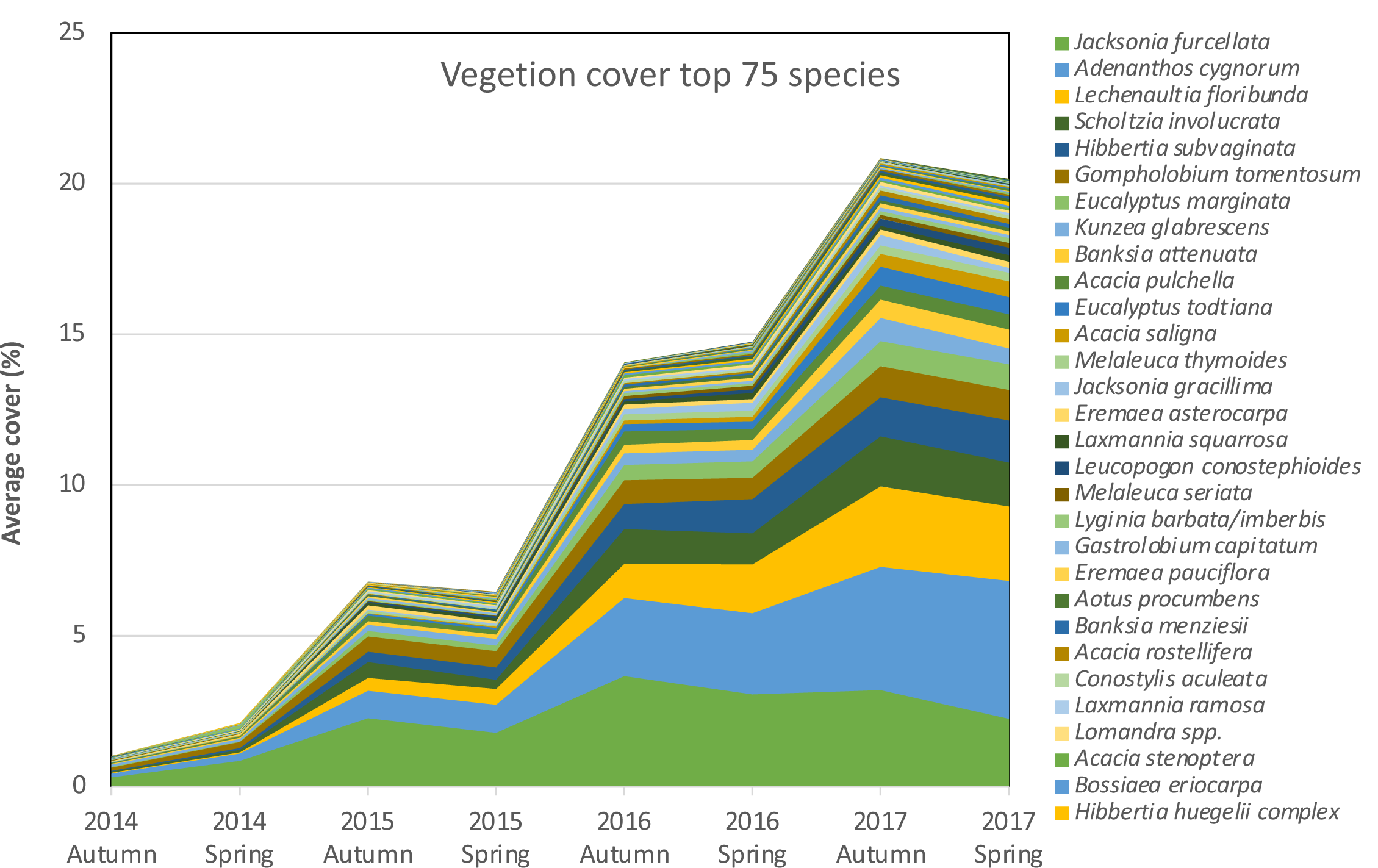
Increases in the average cover of plant species over time in areas with respread topsoil. This graph shows the top 75 species ranked by cover with the 30 most important species listed in order of importance in the legend.

Perennial native plant density peaked at >125,000 stems per ha at AR and ∼100,000 stems per ha at FL in spring 2013. However, most of these were seedlings and many did not survive the severe spring and summer droughts that followed. In the AR topsoil areas, the target density of 7,000 stems per ha was reached by autumn 2015 (Fig. 6). Despite many losses each summer, the average perennial native density increased each year when seedlings emerged each winter. By year 5, there were nearly 43,000 stems per ha of perennial natives (including 23,000 seedlings), far exceeding the target (Fig. 6, Table 2). In contrast, density at FL reached the target in autumn 2015 but declined over the summer of 2016/17, so was below target density in spring 2017, unless seedlings were included (Table 2).

**Figure 6.**
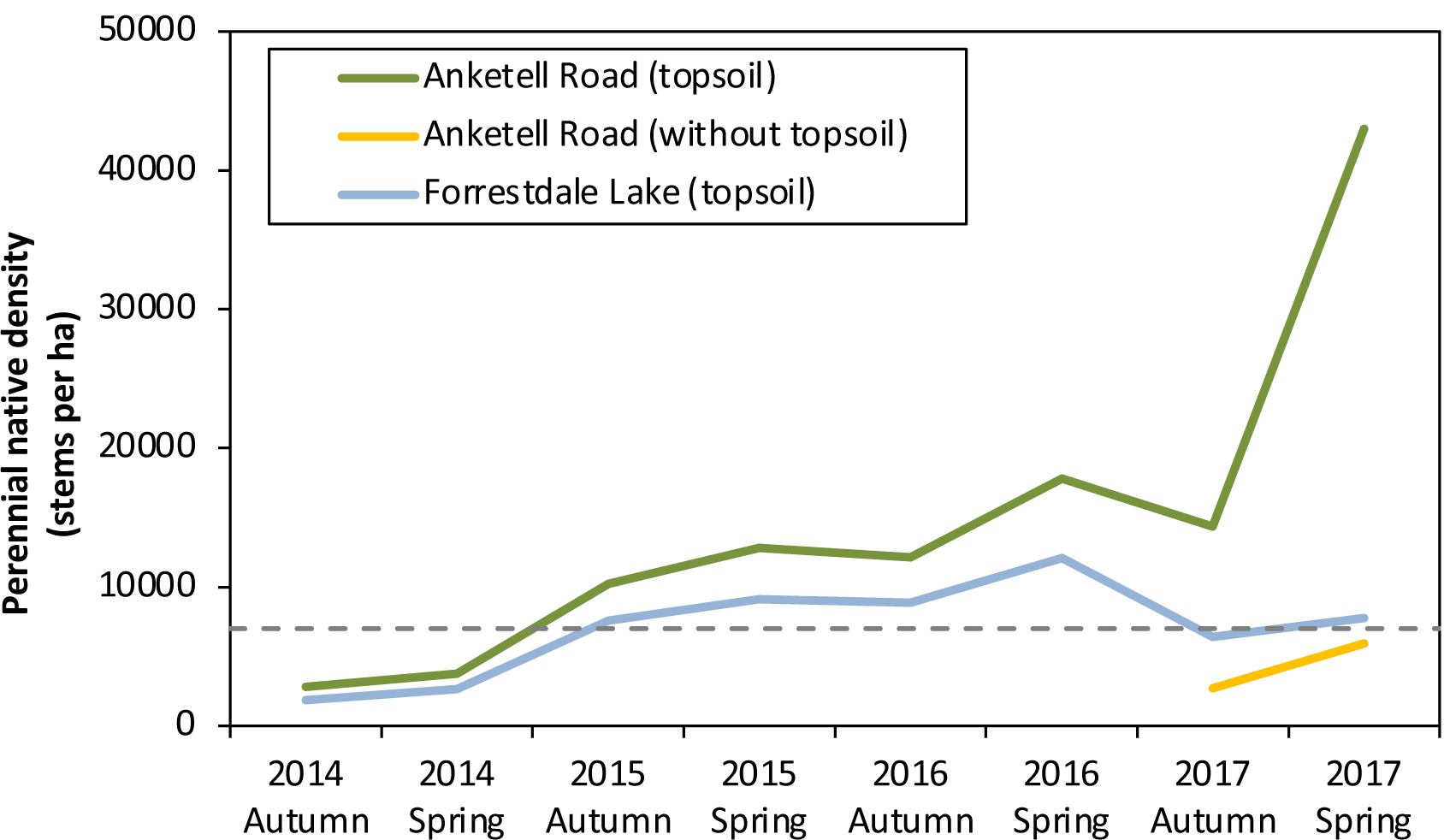
Increases in the average density of perennial native plants in 5×5m quadrats at the Anketell Road restoration site from 2014-2017. The completion criteria target of 7,000 stems per ha is shown by the dashed line. Topsoil areas at Anketell Road are exceeding the target while Forrestdale Lake is just below target (without seedlings). Seedlings were not counted in the 2014 surveys.

Total weed cover, which consisted primarily of annual weeds, peaked at 5% at AR and 6% at FL and has stabilised or declined over time (Fig. 4). Perennial weed density has increased annually since 2014 and fluctuates seasonally more than that of native plants. This is primarily due to the bulbous geophytes *Gladiolus caryophyllaceus, Oxalis pes-caprae, O. purpurea*, and *Romulea rosea*. These plants are small, annually renewing plants, so were a low priority for management. For example, the perennial weeds *Gladiolus caryophyllaceus, Ehrharta calycina, R. rosea* occurred in > 50% of quadrats, but had less than 1% average foliage cover (Fig. 7). Couch grass (*Cynodon dactylon*) had the highest cover at ∼1% in AR areas without respread topsoil. The most abundant perennial weed species were *E. calycina* and *C. dactylon* (Fig. 7). The cover of weeds decreased in most areas due to weed control measures and increasing cover by native plants, but ongoing management will likely be required for perennial grasses.

**Figure 7.**
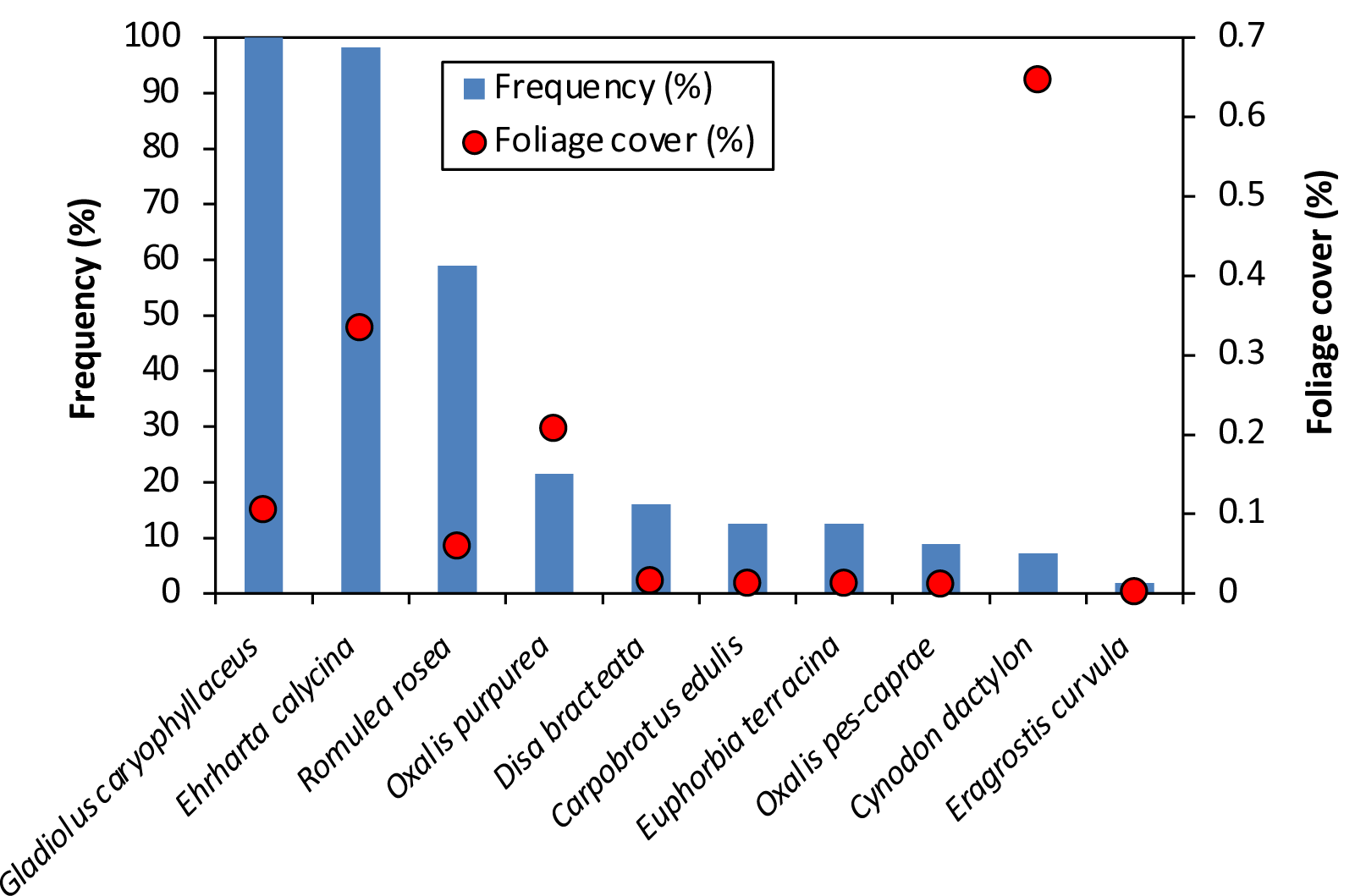
Frequency and foliage cover of perennial weeds at Anketell Road topsoil areas in spring 2017. Some weeds were widespread, but all had low average cover.

Tree density in 25×25 m quadrats at AR reached the 300 stems ha^-1^ target by 2014 in areas where topsoil was spread due to planting and seeding but dropped to 243 by 2017 (Fig. 8). Tree density was lowest at FL (183 stems ha^-1^) and highest at AR areas without applied topsoil (362 stems ha^-1^, Table 2). The latter were in lower-lying areas of the site (dampland ecotones), which may explain higher seedling survival in these areas. Survival of planted tubestock was very low in unfenced areas due to drought and grazing, despite using tree guards. However, most areas had sufficient surviving trees by 2017. Tree diversity was also higher in the non-topsoil areas, with 10 tree species established compared to seven in the topsoil areas, due to inclusion of additional *Melaleuca* and *Eucalyptus* species that grow in dampland ecotone areas.

**Figure 8.**
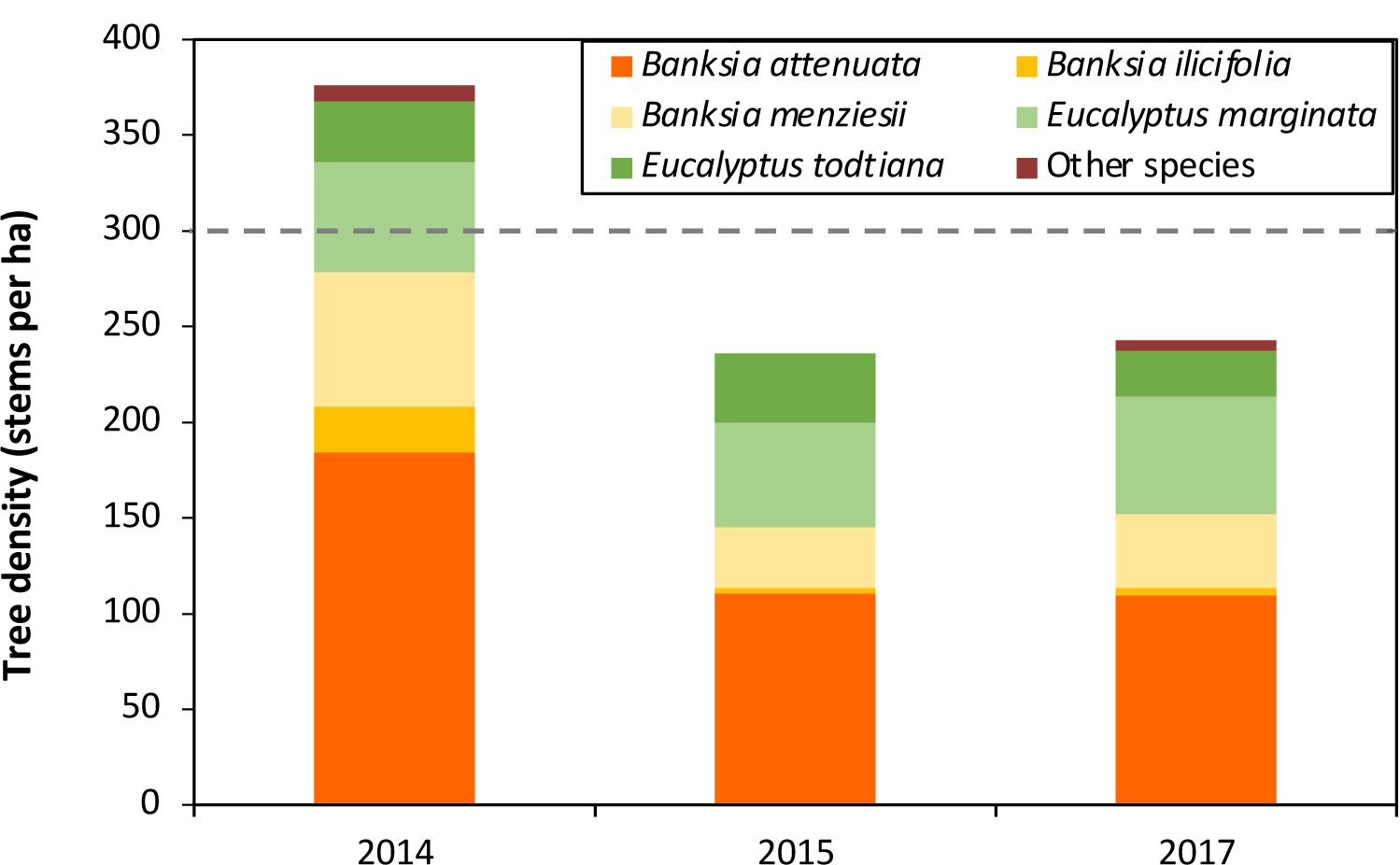
Tree density (stems per ha) from surveys of 25×25 m quadrats at Anketell Road topsoil areas. Tree density exceeded the target of 300 stems per ha (dashed line) in 2014,but has decreased due to drought deaths over exceptionally hot, dry summers.

Differences in growth rates were evident between tree species (Fig. 9), with eucalypt trees increasing by an average of 58 cm per year, while *Banksia attenuata* grew about 50 cm per year and *Banksia menziesii* was slowest at 36 cm per year. However, the latter started flowering when only 3 or 4 years old, so invested more resources in reproduction than other young trees (in 2013 1/3 of *B. menziesii* trees flowered, but <1% of *B. attenuata*). Most five-year-old trees were between 1-2.5 m tall (Fig. 9), but a few were up to 5 m tall.

**Figure 9.**
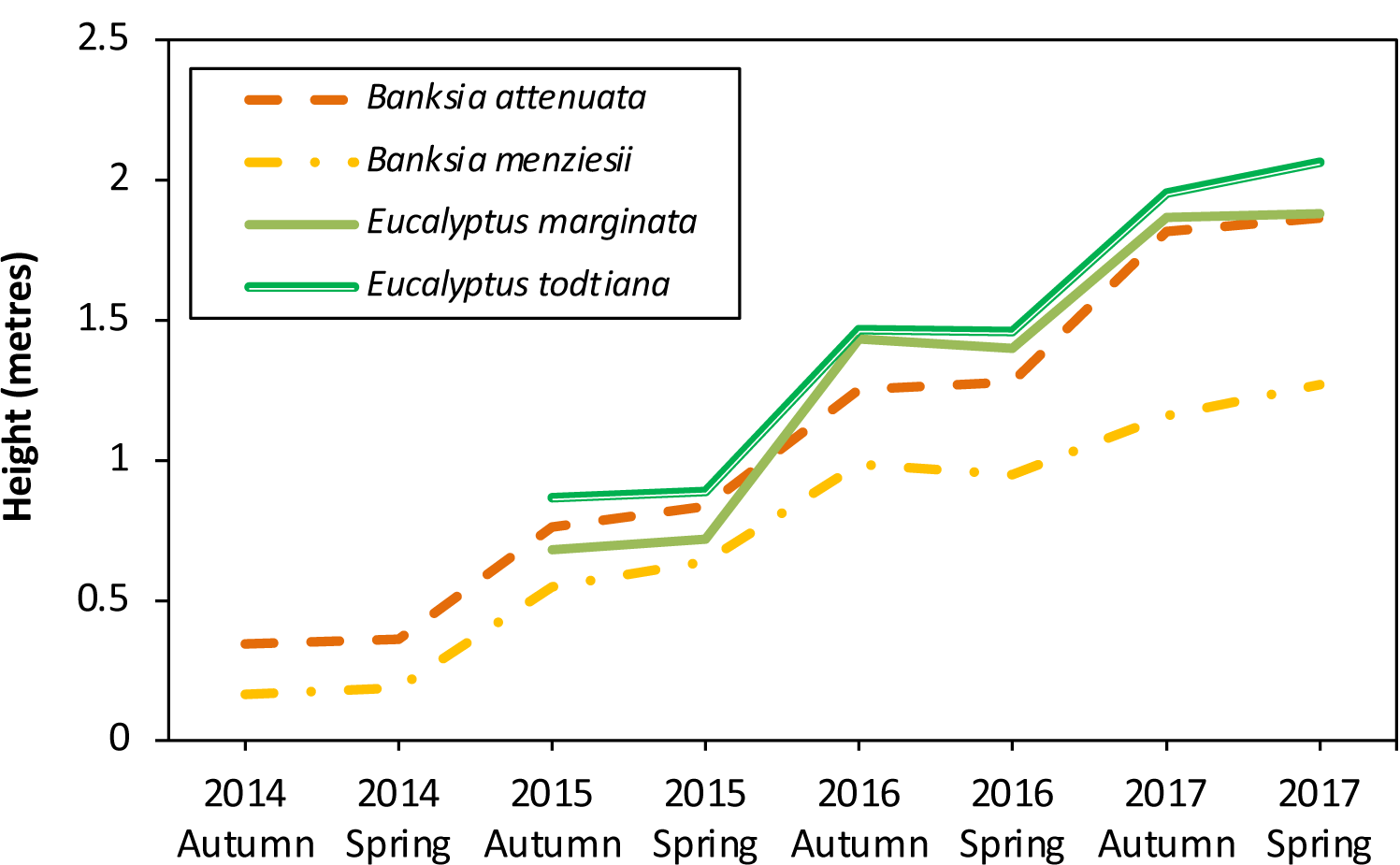
The mean height of dominant tree species in 5×5m monitoring quadrats plotted separately over the first 4 years at Anketell Road topsoil areas. There were two measurements each year in spring (S) and autumn (A). Eucalypt species were not measured in 2014.

After five years, the total diversity of native plants at both sites was similar to topsoil source reference quadrats, with about 162 native plants (Brundrett et al 2018, Fig. 10). The average diversity of plants in 100m^2^ quadrat equivalents reached targets within five years (Table 2). However, there were more uncommon species in the restoration areas than in the reference sites. Most native species found at the topsoil source site either germinated from topsoil, were planted, or were seeded at the restoration sites (Fig. 11). However, there were 10 native species from references sites that were rare in restored areas (Fig. 12) such as, *Hibbertia hypericoides*, a common banksia woodland species. In contrast, 16 native plant species recruited in restoration sites that had not been observed in the topsoil source area (Fig. 12). Some of these spread from adjoining areas, but most were short-lived disturbance opportunists that germinate from topsoil after fire or other disturbances (Fig. 11). The most common of these were annual species of *Austrostipa, Podotheca* and *Trachymene* and the small shrubs *Gastrolobium capitatum* and *Gompholobium tomentosum*. These were initially abundant but were decreasing in numbers by the third year. Common large shrubs (1-3 m high), including *Adenanthos cygnorum* and *Jacksonia furcellata*, became over-dominant in some areas. However, many plants of *J. furcellata* were declining or dead by year 5, with a few new seedlings of this species also emerging (Figs. 5, 13B, G). Disturbance opportunists are expected to continue to decline in areas with developing tree canopies. They have also shed many seeds into the topsoil seed bank which may gradually germinate over subsequent decades. Close to 100 species of weeds were observed in restoration areas. The majority of these are of limited concern, because they are small shade-intolerant annuals that diminished in importance over time as total foliage cover of native plants increased (Fig. 4).

**Figure 10.**
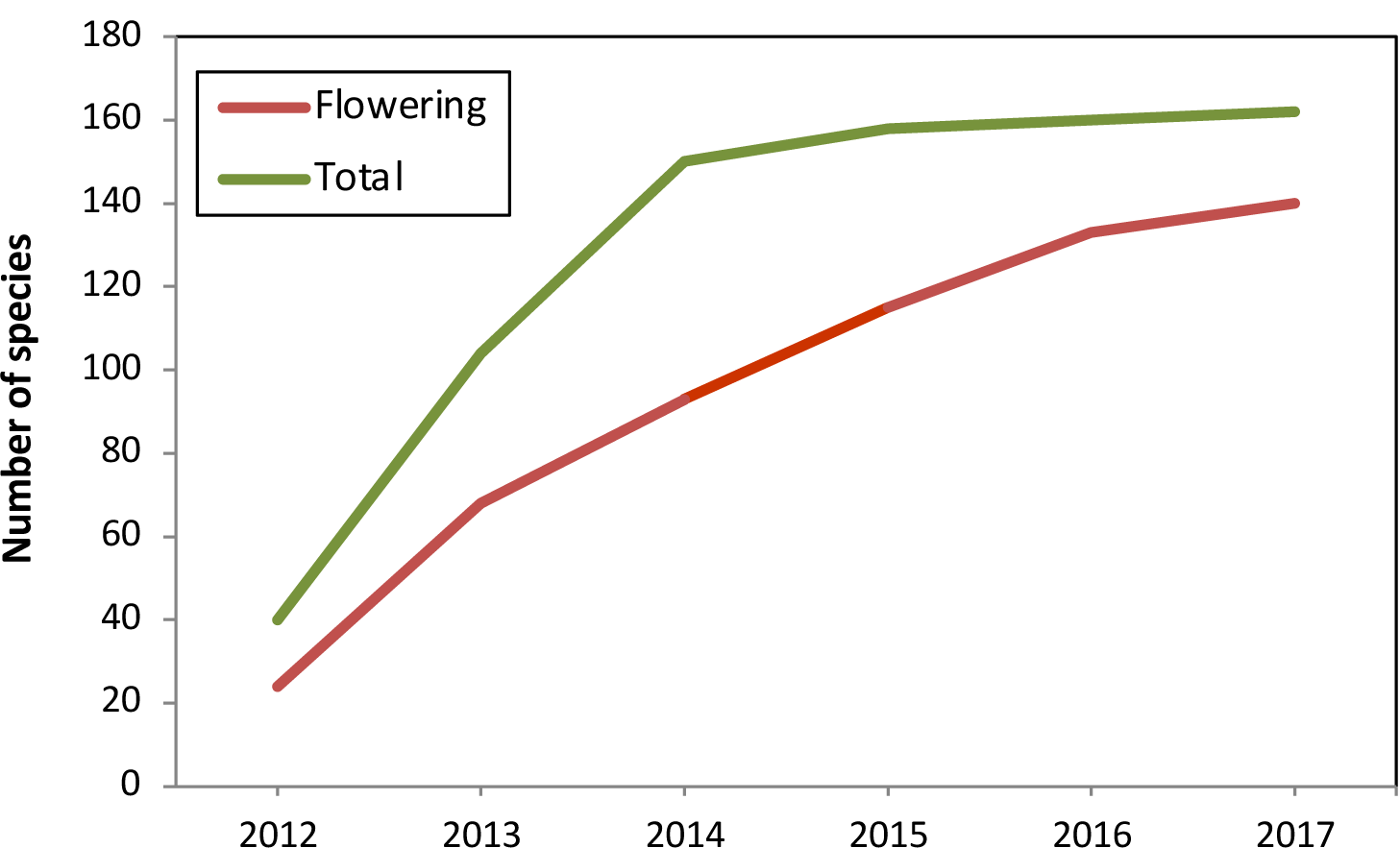
Increases in the number of plant species present and flowering at both restoration sites over the first six years after topsoil transfer. The totals also include species that were planted or direct seeded. Over 86 percent of plants observed have flowered within five years of the initiation of restoration. All species are listed in Table S1.

**Figure 11.**
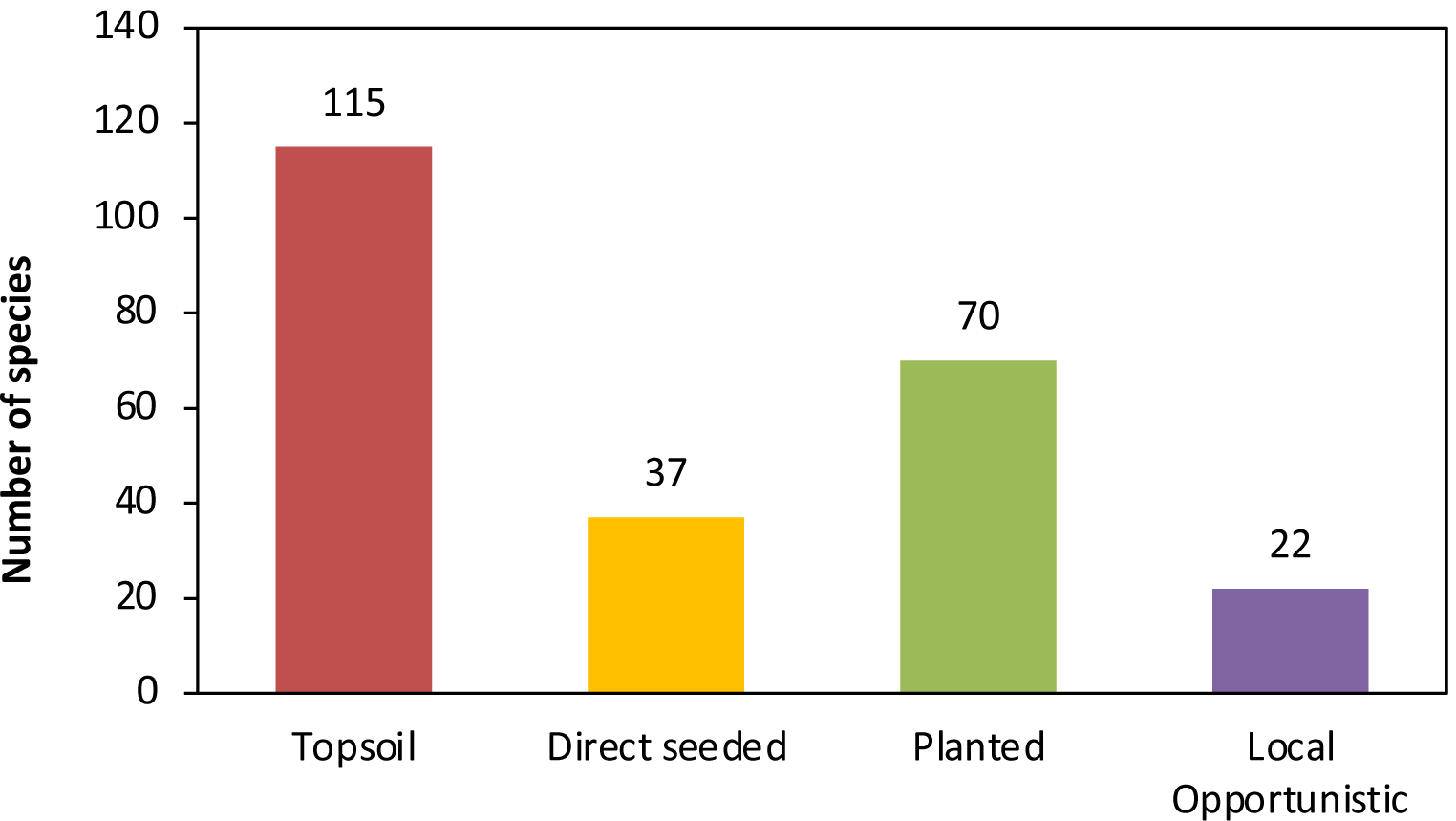
Comparison of the number of native plant species recruited from different propagule sources. There is some overlap between categories, especially for planted and direct seeded species. In total 162 species have been identified in the restoration areas. A complete species list is provided in Table S1.

**Figure 12.**
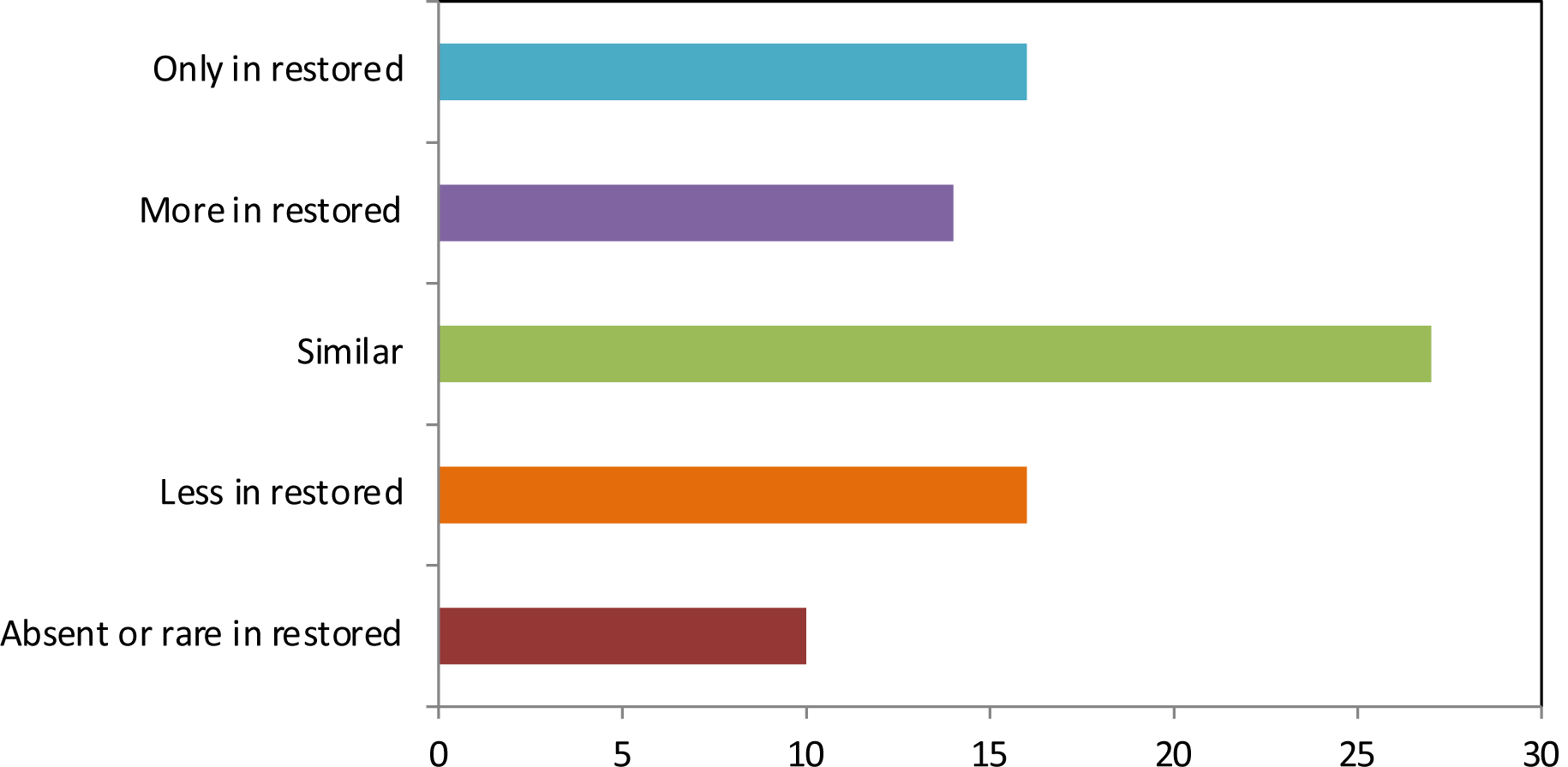
The relative success of species in restored areas relative to the reference site (the topsoil source area) measured as plant occurrence and density.

**Figure 13.**
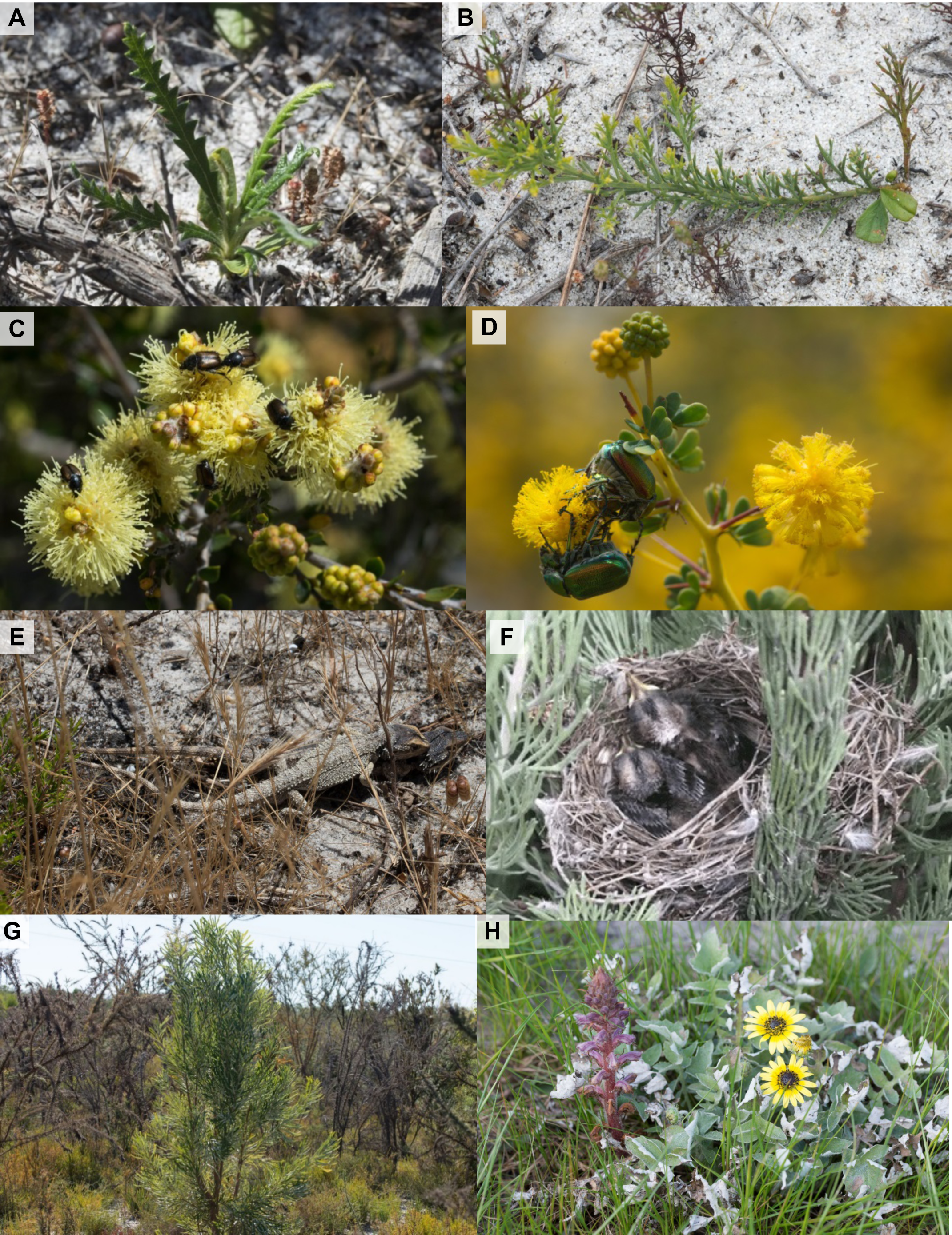
Ecological indicators of restoration success: **A**. Self-sown banksia seedlings found in 2016. **B**. Second generation seedlings of *Jacksonia furcellata* became common as older individuals senesced. **C**. Pollinators, such as nectar scarabs (*Neophyllotocu*s sp. on *Melaleuca thymoides*) were common in spring. **D**. Green spring beetles (*Diphucephala edwardsii* on *Acacia pulchella*) were abundant in winter. **E**. Mating pair of western bearded dragons (*Pogona minor*) in the restoration site. **F**. New Holland honeyeater (*Phylidonyris novaehollandiae*) nestlings in the Anketell Road restoration site. **G**. Declining *Jacksonia furcellata* next to a rapidly growing *Banksia attenuata* tree. **H**. Increased abundance of the parasitic weed *Orobanche* sp. was often linked with *Arctotheca calendula* (Capeweed) decline.

### 2. Spatial variability in restoration sites

After 5 years, 70% of quadrats at AR and 38% at FL had reached the completion criteria target for perennial native plant density (Fig. 14). In areas that received topsoil, plant density was very good or adequate in most areas, but plant density and diversity was typically lower where only seeding and planting occurred (Fig. 15A). Native plant densities in the later occurred because restoration started 2 years later and there was a 1.5 m spacing between seeding rows. Potential reasons for some areas with topsoil with low cover (Fig. 15B) include localised variations in water availability, weed competition, or soil compaction. Areas at FL with poor vegetation establishment tended to coincide with the former locations of buildings, tracks, and market gardens visible in 70-year old aerial photography. These structures were removed after purchase of the land for conservation but have caused localised problems with soil structure or chemistry. In addition, the FL site had a partial canopy of large trees that would strongly compete for water and other resources, especially in summer.

**Figure 14.**
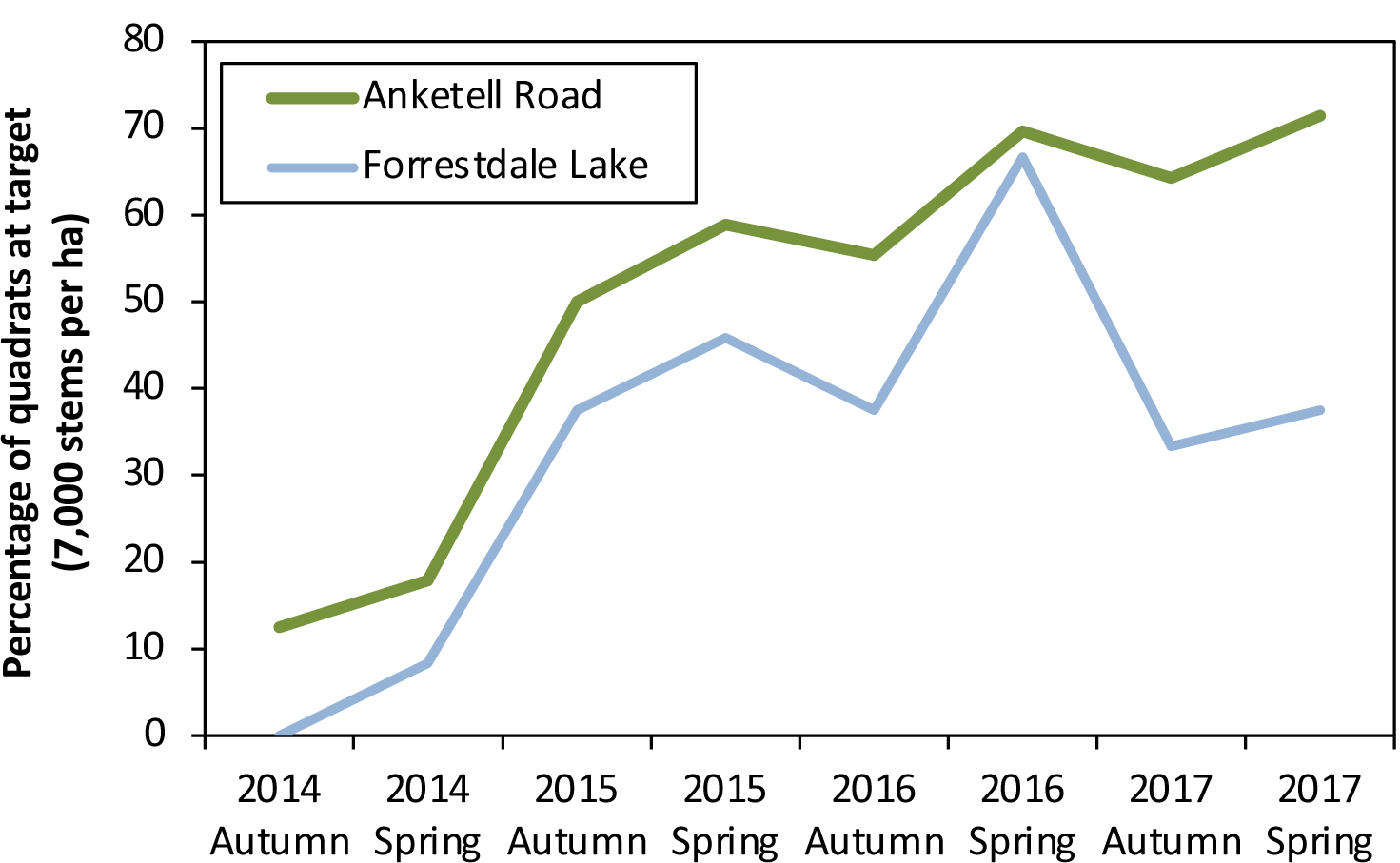
Temporal changes in the percentage of monitoring quadrats reaching target density for perennial natives (7,000 stems per ha) at Anketell Road and Forrestdale Lake (areas restored with topsoil only). Data are from 5×5 m quadrats (n = 56).

**Figure 15.**
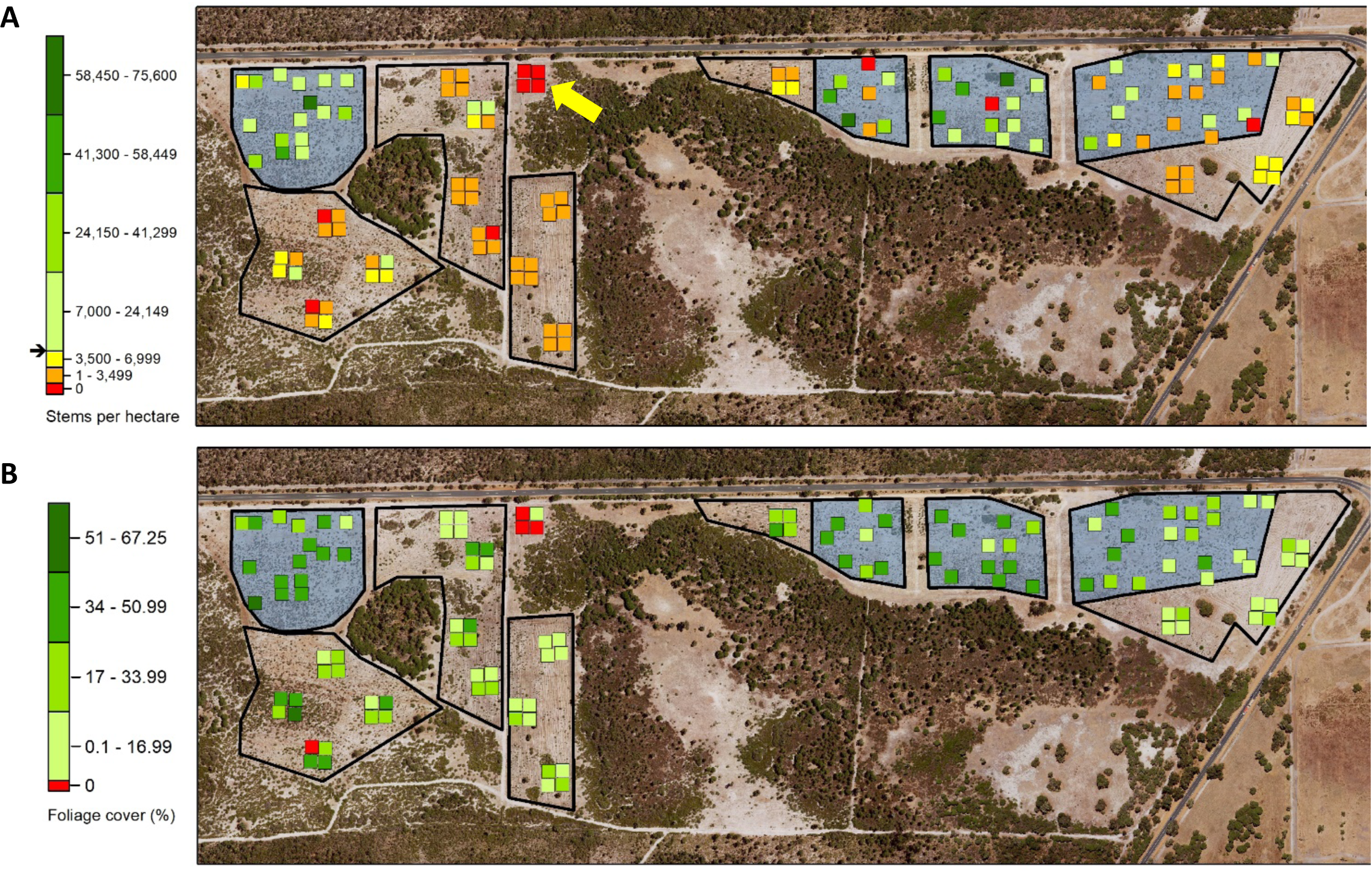
Spatial summaries of restoration outcomes at the Anketell Road restoration site using data from monitoring plots (in 5×5 m quadrats) displayed on an aerial photograph (spring 2017). **A**. Variations in native perennial plant density. Green-shaded quadrats are above the target density of 7,000 stems per ha as indicated on the colour scale. **B**. Foliage cover of native perennials (red = no cover, deeper green colour indicates higher cover). Areas restored without respread topsoil are included for comparison. Blue shading indicates areas are restored with topsoil and black outlines show fencing. A control area without fencing or planting is indicated by yellow arrow. Quadrats are not drawn to scale.

The spatial visualisation of restoration targets allowed more effective management of sites by identifying target areas for infill planting or seeding (Fig. 15). We also found that photo-monitoring at fixed points (Fig. 1) was especially valuable for illustrating changes in vegetation structure over time and identifying variations in restoration outcomes. As an example, Figure 2F shows adjacent areas at AR with good (background) or poor (foreground) vegetation establishment.

### 3. Comparison of methods and sites

The topsoil seed bank provides about 2/3 of the species present at AR and FL restoration areas (Fig. 11). However, this propagule source did not include trees or most large shrubs, as these had canopy-stored seed. In particular, banksia trees were rarely observed germinating in topsoil. This resulted in the need for several complementary methods to achieve adequate outcomes. We also commissioned clonal propagation of selected plant species that were rare or missing from other sources. Some geophytes that recovered quickly apparently grew from tubers rather than seeds transferred in topsoil (Fig 2G).

Figure 16A compares the diversity of native plants that recruited from respread topsoil relative to those primarily from planting and seeding. Lower native species richness was expected in areas without topsoil, due to the absence of species for which seed sourcing and/or germination was not possible. The difference in species richness is most noticeable in the understory, especially for grasses, sedges, and herbs (Fig 16A). Furthermore, results of direct seeding were often very uneven, with some areas of very dense seedlings and other areas where seedlings were widely spaced. However, we were able to fill in large gaps by a second application of seed in 2016 using machine direct seeding or hand seeding in open areas. It was possible to drive around established plants to avoid damaging them when reseeding large areas.

**Figure 16.**
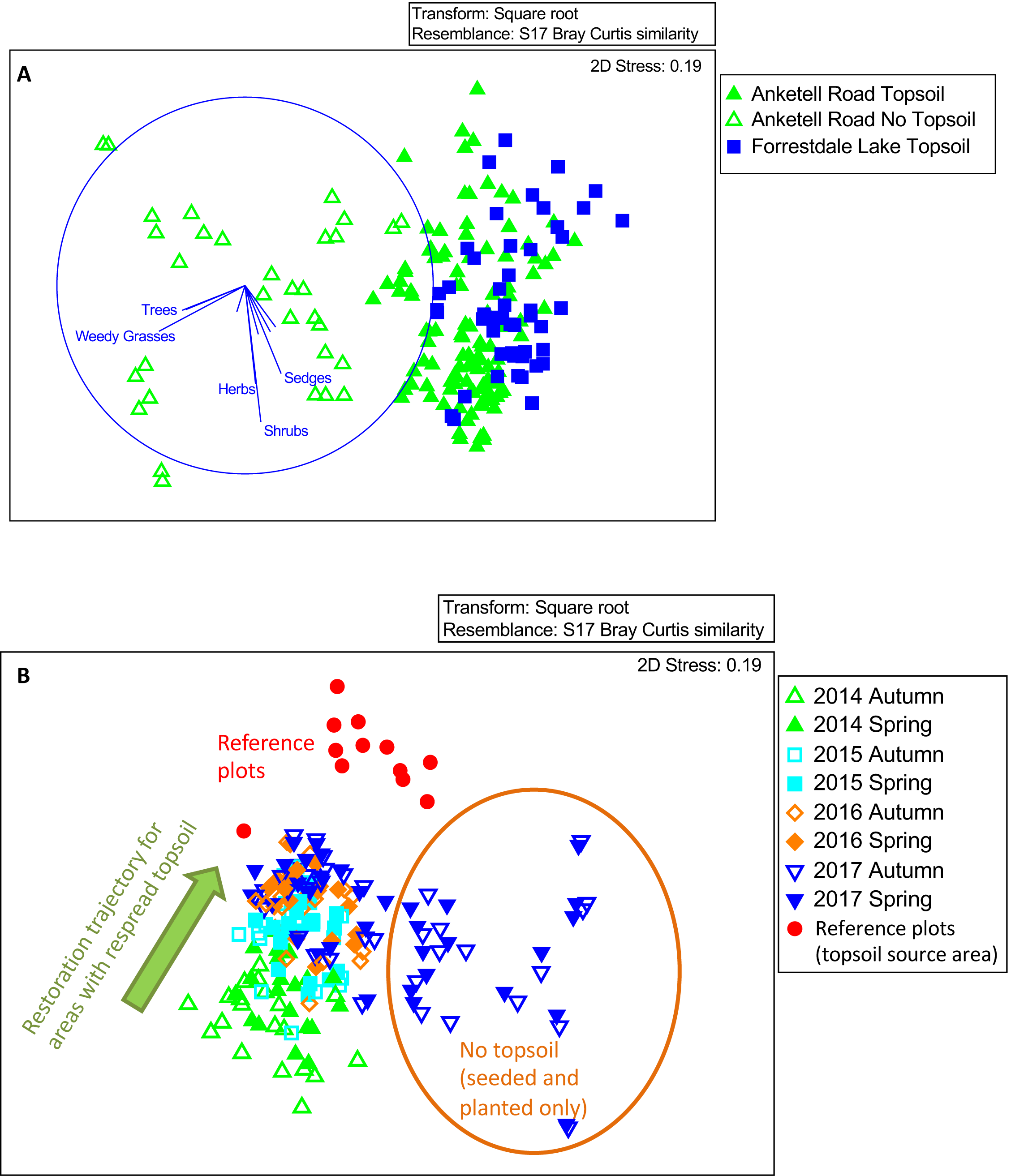
MSD plots showing the cover of perennial plants at the restoration sites. **A**. The floristic composition of quadrats established with respread topsoil, direct seeding, and planting, compared with areas established by direct seeding and planting only, using cover data for species. **B**. Temporal progression in the relative dominance (cover) of native plant species in areas with and without respread topsoil relative to topsoil source reference quadrats (red dots).

Figures 16B and 17 show how the floristic composition and structure of vegetation in restored areas is substantially different from the topsoil source area and is likely to remain so for some time. In general, restored vegetation is less structurally complex with fewer smaller plants, especially sedges and geophytes. It is expected that differences between restored and reference sites will decrease over time as native plants grow and spread, weeds are supressed by shading and plants disperse within sites. We observed a turnover of native plants by year 5 as disturbance opportunists were gradually replaced by long-lived species, some of which expand very slowly due to clonal growth. Figure 17 also shows major differences in vegetation structure between reference sites that were a few km apart. Defining differences between restored areas and reference sites is made more complicated by the very high variability of structural and taxonomic diversity within all banksia woodland habitats.

**Figure 17.**
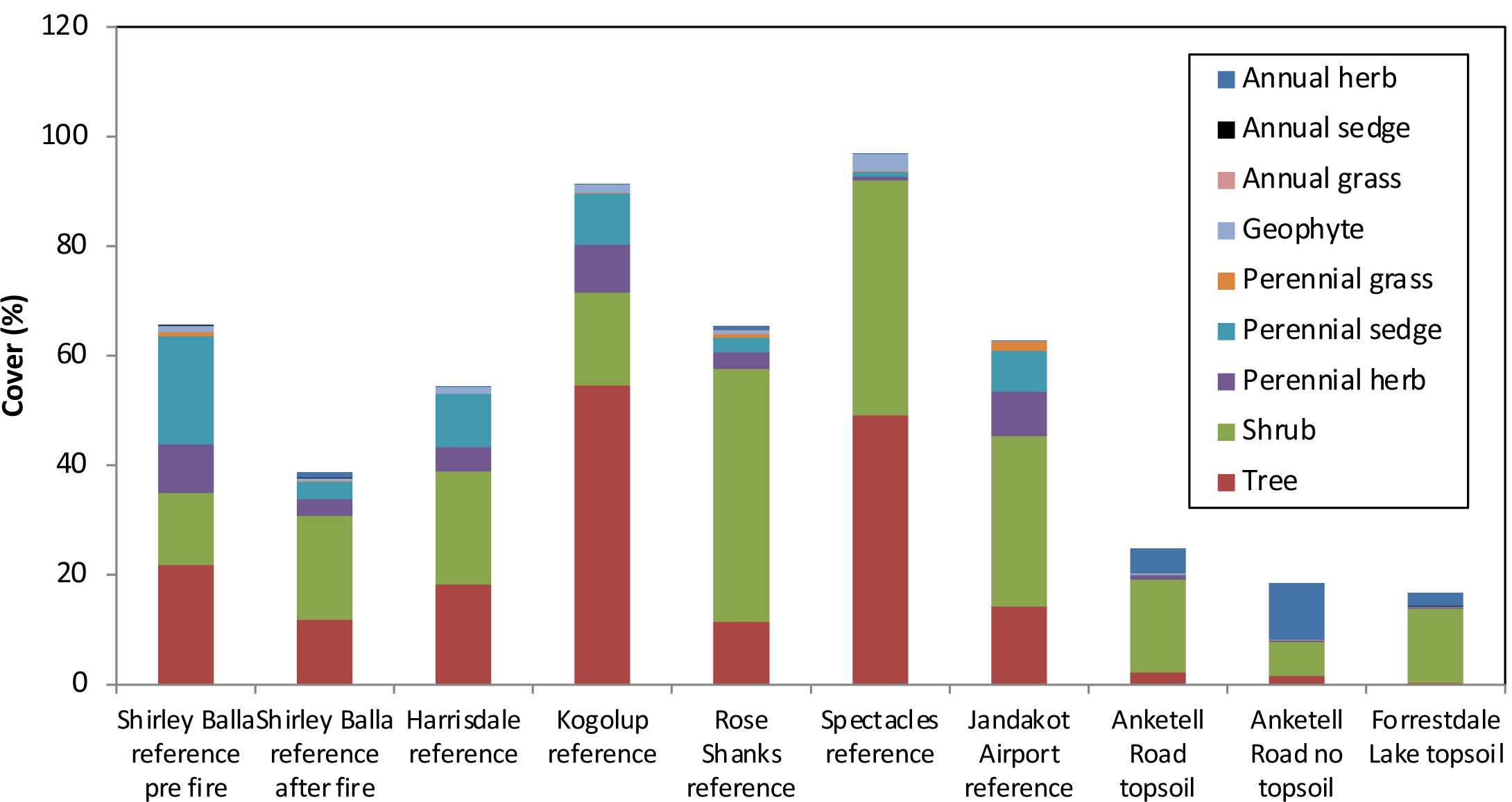
Cover of native plants by growth form for six reference sites (including one for pre and post fire condition) and restoration sites reveals vegetation structure varied considerably between reference sites, between reference sites and restored areas (last 3 columns), and between restored areas.

### 4. Restoration outcomes relative to targets

After five years, most of the completion criteria targets in Table 2 had been reached or exceeded. However, seed germination from the topsoil and the survival of seedlings was highly variable across sites so there were some areas with insufficient native plant density and cover. The same restoration methods at FL resulted in lower density of perennial natives and trees than at AR (Table 2). This is likely due to historical land management impacts on soil properties, as explained above. Areas restored with seeding or planting only were not expected to reach the same native plant density as areas with topsoil in the short term, but tree density had exceeded targets within 3 years. This approach, which was much less expensive than topsoil transfer, was very effective for establishing trees and shrubs, including the most important food plants for Carnaby’s cockatoo (Table 2). Some additional planting or seeding may be required to reach targets for tree density, especially at FL. However, by year 5 trees were producing abundant seed so dispersal and germination within sites should be sufficient to eventually reach targets. It will be necessary to monitor future vegetation development and recruitment within sites to determine if any additional management actions are required.

### 5. Recovery of ecological interactions

Over 86% of native species flowered within five years of the initiation of restoration activities (Fig. 10). Long-lived perennial plants, especially trees, required several years longer to flower than most shrubs and herbs. Substantial pollinator activity by a wide diversity of native insects and abundant European honeybees was observed on site (see video Figure S1). This resulted in abundant seed set at both restoration sites. For trees, flowering of *Banksia menziesii* commenced in 2015, *B. attenuata* and *Eucalyptus marginata* in 2016, and *Melaleuca rhaphiophylla* and *M. preissiana* in 2017. Other plants that have flowered prolifically include species of *Jacksonia, Lechenaultia, Melaleuca* and *Kunzea*, as well as the native orchid *Caladenia flava* (Fig. 2G).

As explained above, we observed early stages of plant succession in restored areas due to the decline of some species which germinated abundantly from topsoil in the first 3 years. These included *Hibbertia subvaginata, Jacksonia furcellata* and *Gompholobium tomentosum*, which are also abundant after fire in banksia woodland. These three species matured early, flowered prolifically so should have already produced sufficient seed to replenish the soil seed bank. Declining cover by *J. furcellata* was linked to increasing dominance by trees in some areas, so may result from competition for water (Fig. 13G). We also observed second generation seedlings of banksia species (Fig. 13A) and *J. furcellata* (Fig 13B).

After 5 years there is increased evidence of kangaroo activity within fenced areas (scats and grazing have been detected), probably because the plastic sight lines used to deter them have become less effective over time (Fig. 2H). However, these fences provided adequate protection during the early establishment of vegetation. Other examples of harmful or beneficial ecological interactions observed at restoration sites that provide evidence of ecosystem complexity and sustainability we observed include:

1. Grazing of banksia seedlings by invertebrates.
2. Broomrape (*Orobanche minor*) was a common parasitic plant in restored areas but primarily attacked weed species. This resulted in reduced vigour of cape weed (*Arctotheca calendula*) in some areas (Fig 13H).
3. Mycorrhizal and saprophytic fungi were observed to fruit in restored areas within a few years.
4. Parasitic galls caused substantial impacts to some plants, especially *Acacia* spp.
5. The smut fungus (*Tilletia ehrhartae*) is attacking the weedy grass *Ehrharta calycina*.
6. Mating of reptiles such as the western bearded dragon (*Pogona minor*) (Fig. 13E).
7. Nesting of birds such as New Holland honeyeaters (*Phylidonyris novaehollandiae*) (Fig. 13F).

## Discussion

One of the key objectives of this project was to evaluate the relative cost-effectiveness of different methods for restoration of banksia woodland (preliminary costings are provided in Brundrett et al. 2015). Topsoil transfer was the most efficient method for restoring native plant diversity, but supplementary planting or seeding was also required to establish species that were absent from the topsoil seed bank. It also needs to be acknowledged that topsoil transfer has not always been successful in other restoration projects, since it requires topsoil source areas to be free from major weeds, diseases such as *Phytophthora* dieback and to contain sufficient amounts of viable seed. The harvesting, transfer and respreading of the topsoil also needs to be carefully planned and executed for successful results, since topsoil must be dry, only stored briefly, not mixed with subsoil and should be transferred in autumn (Rokich et al. 2000, Rokich 2016).

The germination of West Australian plants from the topsoil seed bank have been the focus of previous studies that investigated their diversity and factors that promote their germination. These include the impacts of heat and smoke on seed germination (Bell 2001, Meney et al. 1994, Roche et al. 1997, Rokich & Dixon 2007). Fowler *et al*. (2015) measured germination in the glasshouse using topsoil from the same location used in this study (Jandakot Airport) and Waryszak et al. (in press) measured recruitment over 2012 to 2014 at the same restoration sites that we used. Both studies confirmed that severe disturbance during topsoil transfer promoted abundant seed germination for a subset of banksia woodland plants in a similar way to fire or smoke. The later study also found that additional treatments to promote germination or seedling survival had little further impact and corroborated our observations of very high seedling mortality over summer (Waryszak *et al*. in press). In the current study our main observations concerning plant diversity outcomes after topsoil transfer were:

1. The majority of species were either much more or much less common in restored areas relative to reference sites and a few species were only present in either restored or reference sites.
2. Plant species that were disturbance opportunists were much more abundant in young restoration sites relative to reference sites (arising from the soil seed bank). These species started to decrease after a few years, but set many seed so are expected to remain dormant in the soil seed bank until the next disturbance.
3. We observed that our restoration sites were initially more like banksia woodland that is recovering after fire, as many of the same disturbance opportunists are initially common in both systems.
4. Many long-lived species were very slow to become established, especially those with clonal growth and those that resprout vigorously after fire.
5. Weeds and annual native plants are initially very abundant but declined over time (the former due to appropriate weed management).
6. Tree cover and density was very slow to recover and banksias were more difficult to establish than eucalypts.

In general, direct seeding using seed drill technology (e.g. Jonson 2010) was very efficient and successful at restoring banksia woodland plants, but plant diversity and density was substantially lower than in areas which also had respread topsoil. In all cases, we found that several years of planting and seeding were required to reach targets because of high summer attrition. We also noted there were issues with site properties that impacted on plant establishment, even after a rigorous site selection process (Brundrett et al. 2019). In particular, we found that areas with the most intensive historic land use (horticulture) were less suitable for restoration than areas with less intensive use (grazing).

To achieve restoration targets, our project used an adaptive management framework with progress determined by annual monitoring (Brundrett et al. submitted). This required low survival rates or difficulty establishing certain species to be counteracted by infill planting and seeding on subsequent years. Adaptive approaches were also used to control threats such as weeds and grazing and experimental trials used to compare the effectiveness of different restoration methods. Other examples included direct seeding and planting of additional seedlings where it was needed most and targeting perennial weeds which had cover values that increased most rapidly. Restoration monitoring data was also used to help adjust seed collecting and nursery orders. However, species available for planting and seeding was restricted by what could be obtained by seed collectors and could also be successfully germinated or grown by nurseries. Some of the species that were most common at Jandakot Airport bushland such as the sedge-like plants *Desmocladus flexuosus* and *Lyginia barbata* are not easily grown from seed, so we used vegetative propagation for these species.

Seed collecting was a major expense and challenging for many species, it is most effective for plants with canopy stored seed and/or abundant synchronised fruiting. Fortunately, many of the species that failed to grow from topsoil were amenable to seed collection and production in nurseries (see Fig. 11). However, these species varied considerably in their mortality due to summer drought and many had issues with poor seed quality and/or germination that we needed to overcome by seed science research (this will be the subject of a subsequent paper).

The structure and composition of a recently disturbed habitat should be similar to reference ecosystems and contain all the key species required for eventual recovery of vegetation structure and function. (Environmental Protection Authority 2006, McDonald et al. 2016). Restored banksia woodland is unlikely to have similar floristic composition or vegetation structure to reference sites in the short term. These differences arise because the plant functional traits that determine their capacity to colonise new habitats vary considerably between species (Lamont et al. 1991, Bellairs & Bell 1993, Rokich & Dixon 2007, Tozer et al. 2012, Keith 2012). For example, there are a substantial number of species which rarely germinate from topsoil, did not produce readily collectible seed, and/or could not be readily propagated in nurseries, so are usually rare or absent in restoration sites.

Restoration in a completely degraded area represents a secondary ecological succession rather than a primary succession if the site has existing vegetation or propagules. The classical view of secondary ecological succession is that, provided environmental conditions remain the same, vegetation will develop through a series of floristic communities towards a single ‘climax’ vegetation type, a process described as “Relay Floristics” (Egler 1954). However, previous mine site restoration studies have found that while vegetation structure appears to progress readily towards a pre-mining state, the floristic community composition is highly influenced by the species that establish first and that this composition is difficult to alter after it has established (Norman et al. 2006, Koch 2007). This means that it is important to ensure the initial species diversity is close to that required in the final restored banksia woodland. There is a fine line between setting sufficiently rigorous completion criteria to ensure sufficient outcomes while avoiding criteria which cannot be attained due to operational constraints. In cases where adequate restoration of the original plant communities is unlikely to be possible due to operational constraints, this should be acknowledged during the project approval process and actions that are more effective in mitigating environmental impacts should be considered (Environmental Protection Authority 2006).

Completion criteria are designed to answer a number of key questions about the structure, composition, and sustainability of ecosystems relative to reference conditions (McDonald et al. 2016). They are also required to ensure sufficient cover to stabilise the site, inhibit weed growth and provide visual amenity (Environmental Protection Authority 2006). However, we could not find many examples of their effective use for banksia woodland restoration. In particular, it was difficult to find examples of projects which provide data on tree density (especially for banksias), or data on other keystone species, or on vegetation structure. It is also uncommon to find meaningful plant diversity comparisons with reference sites (using the same quadrat areas) or floristic community type comparisons (see Brundrett et al. submitted). As explained in the introduction, it is essential to set more effective targets for banksia woodland restoration projects in the future so that doubts about the environmental benefits of these offset-funded projects can be addressed. In particular, completion criteria that measure floristic composition relative to appropriate local plant communities. It is also necessary to measure the establishment of keystone species such as trees separately. In our sites, ecosystem diversity was also maintained by creating separate species lists for upland and ecotone dampland habitats based on appropriate reference sites.

Restoration projects require adaptive management actions informed by comprehensive monitoring to ensure that species are returning to density, cover and frequency values found in reference habitats (Environmental Protection Authority 2006, McDonald et al. 2016). This requires reporting of outcomes relative to completion criteria, but some of these are often arbitrary. Plant diversity totals can be of limited value, as species detected will largely be determined by sampling effort and it is much easier to detect uncommon species in relatively open areas (Brundrett et al. submitted). At our sites, a 60% plant diversity target relative to the reference site equated to the proportion of species detected by monitoring sufficient quadrats to also measure plant density and cover accurately (Brundrett et al. submitted).

We also suggest that different plant growth stages should be counted separately to monitor restoration outcomes. For example, we observed many more seedlings than older individuals of perennial native plants, but their attrition was exceedingly high over summer. The short-term nature of this project made establishment of cover values similar to those of undisturbed banksia woodland unachievable, but similar plant densities and diversity were measured. We expect that several decades of monitoring will be required to establish realistic completion criteria for areas restored by seeding or planting only. Components such as litter, fallen woody debris and habitat for fauna and fungi will also take many years to recover. However, monitoring reproduction (flowering and set seed) can provide a powerful interim measure of regenerative potential and helps to predict resilience after fire.

Our comprehensive restoration monitoring programs resulted in new insights into how measuring restoration progress can inform adaptive management. For example, we found that completion criteria maps were a much more powerful tool for visualising outcomes than average values for entire sites (e.g. Fig. 13). This approach enabled targeted infill planting and helped identify site specific issues that limited plant establishment. The proportion of monitoring quadrats that meet targets is also very effective for summarising outcomes (e.g. Fig. 12). Taking photographs from fixed points several times a year was another powerful tool for restoration management, in combination with downward facing photos, aerial photographs and remote sensing images that can all be efficiently converted into cover values (Brundrett et al. 2018, van Dongen et al. submitted).

Overall, we are confident that banksia woodland can be successfully restored on a large scale, provided that topsoil of sufficient quality can be obtained and there are sufficient resources for seed collecting, planting of seedlings, site management and monitoring. However, the effectiveness of this project was seriously impacted by interrupted funding and other issues with the regulation and enforcement of offsets (see Brundrett et al 2018). We also observed substantial differences in structure and composition between restored and reference habitats that are likely to persist in the short term. Thus, long-term monitoring is required to measure how these differences change over time due to plant dispersal within and into sites.

We also observed that some of the hottest and driest summers on record (www.bom.gov.au/climate/history) resulted exceptionally high seedling mortality in our sites. This requires further research to determine how restoration projects are likely to be impacted by the extreme climatic events (and increased fire frequencies) that are expected in Western Australia in the future (CSIRO and Bureau of Meteorology 2015). We have also become aware that there is a need to share knowledge with members of community groups, industry, and government agencies about what is required for effective restoration of banksia woodland. A database on traits that regulate the restoration potential of plants and how to overcome them is needed. A research centre that investigates how to restore banksia woodland most efficiently and effectively would improve outcomes for all restoration projects and establish better communication for everyone involved.

## Acknowledgements

Funding was provided by Jandakot Airport Holdings under Commonwealth ministerial conditions (EPBC 2009/4796). The project team for 2011-2017 also included Karen Jackson, Sapphire McMullan-Fisher, Julie Fielder, Alice O’Connor, Tanya Llorens, Tracey Moore, Tracy Sonneman, Julia Cullity, Karen Bettink, Brett Glossop, and Mathew Woods. We thank all the people who helped with planting including DBCA staff, Green Skills Inc., Green Army, Friends of Forrestdale, Birdlife Australia, and others. We also thank volunteers who helped with monitoring. Machine direct seeding was undertaken in collaboration with Greening Australia WA. Work at the DBCA Threatened Flora Seed Centre was managed by Anne Cochrane and Andrew Crawford.

## Supplemental Data

**Table S1**. Native plant species established in restoration sites in the first six years (2012-2017, species total 162), their primary means of recruitment and time to first flowering. Species propagated by clonal division are also shown.

**Table S2**. List of open access reports containing additional information about this project.

**Figure S1**. Video of pollinator activity at restoration sites: https://www.youtube.com/watch?v=SpHgpDwYzH8

**Figure S2**. Video summarising project: https://www.youtube.com/watch?v=xtNdUL4p5Gc&feature=youtu.be

