## Supplementary material for "Several complementary methods of plant establishment are required for effective restoration of banksia woodland": Supplimental Table S1

**Table S1.** Native plant species established in restoration sites in the first six years (2012-2017, species total 162), their primary means of recruitment and time to first flowering. Species where planted seedlings were propagated by clonal division are also indicated (*).

| Species | Topsoil | Topsoil and other | Direct Seeded | Planted 2012 -2016 | Local Opportunist | Dampland ecotone only | First started flowering |
|---|---|---|---|---|---|---|---|
| *Acacia huegelii* | 1 | | | 1 | | | 2014 |
| *Acacia pulchella* | 1 | | | 1 | | | 2014 |
| *Acacia saligna* | 1 | | | 1 | | | 2016 |
| *Acacia stenoptera* | 1 | | | | | | 2015 |
| *Adenanthos cygnorum* | 1 | | | | | | 2015 |
| *Allocasuarina fraseriana* | | | 1 | 1 | | | |
| *Allocasuarina humilis* | | | 1 | 1 | | | 2016 |
| *Amphipogon turbinatus* | 1 | 1 | 1 | 1 | | | 2014 |
| *Anigozanthos humilis* | 1 | 1 | 1 | | | | 2013 |
| *Anigozanthos manglesii* | 1 | 1 | 1 | 1 | | | 2012 |
| *Aotus procumbens* | 1 | | | 1 | 1 | | 2013 |
| *Arnocrinum preissii* | 1 | | | | | | 2013 |
| *Austrostipa compressa* | 1 | 1 | 1 | | 1 | | 2012 |
| *Austrostipa macalpinei* | | | | | 1 | | 2014 |
| *Babingtonia camphorosmae* | | | 1 | | | | 2015 |
| *Banksia attenuata* | rare | | 1 | 1 | | | 2016 |
| *Banksia ilicifolia* | | | 1 | 1 | | | 2016 |
| *Banksia menziesii* | | | 1 | 1 | | | 2015 |
| *Beaufortia elegans* | | | | 1 | | 1 | |
| *Boronia ramosa* | 1 | | | | | | 2013 |
| *Bossiaea eriocarpa* | 1 | 1 | 1 | 1 | | | 2014 |
| *Brachyloma preissii* | 1 | | | 1 | | | 2014 |
| *Burchardia congesta* | 1 | 1 | 1 | | | | 2014 |
| *Caladenia flava* | 1 | | | | | | 2014 |
| *Calandrinia corrigioloides* | 1 | | | | | | 2013 |
| *Calandrinia granulifera* | 1 | | | | | | 2013 |
| *Calothamnus lateralis* | | | | 1 | | 1 | 2017 |
| *Calytrix angulata* | | | | 1 | | | 2013 |
| *Calytrix fraseri* | | | | 1 | | | 2014 |
| *Cartonema philydroides* | 1 | | | | 1 | | 2012 |
| *Cassytha flava* | 1 | | | | 1 | | 2015 |
| *Centrolepis drummondiana* | 1 | | | | | | 2012 |
| *Centrolepis inconspicua* | 1 | | | | | | 2013 |
| *Chamaescilla corymbosa* | 1 | | | | | | 2013 |
| *Comesperma calymega* | 1 | | | | | | 2013 |
| *Conostephium pendulum* | 1 | | | | | | 2016 |
| *Conostylis aculeata* | | | 1 | 1 | | | 2014 |
| *Conostylis juncea* | 1 | | | | | | 2014 |
| *Conostylis setigera* | 1 | | | 1 | | | 2014 |
| *Corymbia calophylla* | | | 1 | 1 | | 1 | |
| *Corynotheca micrantha* | 1 | | | | | | 2015 |
| *Crassula colorata* | 1 | | | | | | 2012 |
| *Crassula decumbens* | 1 | | | | | | 2013 |
| *Croninia kingiana* | 1 | | | | | | 2014 |
| *Dampiera linearis** | 1 | | | 1 | | | 2013 |
| *Dasypogon bromeliifolius* | 1 | 1 | 1 | 1 | | | 2013 |
| *Daucus glochidiatus* | 1 | | | | | | 2014 |
| *Daviesia physodes* | 1 | | | | | | 2016 |
| *Daviesia triflora* | 1 | | | | | | 2017 |
| *Desmocladus flexuosus* | 1 | | | 1 | | | |
| *Dianella revoluta* | | | | 1 | | | |
| *Dichopogon capillipes* | | | | 1 | | | 2017 |
| *Diuris corymbosa* | 1 | | | | | | 2014 |
| *Drosera erythrorhiza* | 1 | | | | | | 2016 |
| *Drosera glanduligera* | 1 | | | | | | 2012 |

| Species | Topsoil | Topsoil and other | Direct Seeded | Planted 2012 -2016 | Local Opportunist | Dampland ecotone only | First started flowering |
|---|---|---|---|---|---|---|---|
| *Drosera macrantha* | 1 | | | | | | 2013 |
| *Drosera paleacea* | 1 | | | | | | 2015 |
| *Epilobium hirtigerum* | | | | | 1 | | 2013 |
| *Eremaea asterocarpa* | | | 1 | 1 | | | 2013 |
| *Eremaea pauciflora* | 1 | | 1 | 1 | | | 2015 |
| *Eucalyptus marginata* | | | 1 | 1 | | | 2016 |
| *Eucalyptus rudis* | | | | 1 | 1 | 1 | 2017 |
| *Eucalyptus todtiana* | | | 1 | 1 | | | 2017 |
| *Exocarpos sparteus* | | | | | 1 | | 2012 |
| *Gastrolobium capitatum* | 1 | 1 | 1 | | | | 2014 |
| *Gnephosis angianthoides* | 1 | | | | | | 2012 |
| *Gompholobium tomentosum* | 1 | 1 | 1 | 1 | | | 2013 |
| *Gonocarpus pithyoides* | 1 | | | | | | 2013 |
| *Haemodorum spicatum* | 1 | 1 | 1 | | | | 2013 |
| *Hakea prostrata* | | | 1 | | | | 2015 |
| *Hardenbergia comptoniana* | 1 | | | | 1 | | 2016 |
| *Hemiandra pungens* | 1 | | | 1 | 1 | | 2013 |
| *Hemiandra* sp. Jurien | | | | 1 | | | 2016 |
| *Hensmania turbinata* | 1 | | | | | | 2014 |
| *Hibbertia huegelii/sericostachya** | 1 | | 1 | 1 | | | 2013 |
| *Hibbertia hypericoides* | 1 | | | 1 | | | 2013 |
| *Hibbertia racemosa* | | | | 1 | | | 2014 |
| *Hibbertia subvaginata** | 1 | | | 1 | | | 2012 |
| *Homalosciadium homalocarpum* | 1 | | | | | | 2012 |
| *Hovea trisperma* | 1 | | | | | | 2013 |
| *Hyalosperma cotula* | 1 | | | | | | 2013 |
| *Hypocalymma angustifolium* | 1 | 1 | 1 | 1 | | | 2014 |
| *Hypocalymma robustum* | 1 | | | | | | 2014 |
| *Hypolaena exsulca* | 1 | | | | | | 2015 |
| *Isolepis marginata* | 1 | | | | 1 | | 2012 |
| *Jacksonia furcellata* | 1 | 1 | 1 | 1 | 1 | | 2014 |
| *Jacksonia gracillima* | 1 | | | | | | 2015 |
| *Jacksonia sternbergiana* | 1 | | | | | | 2014 |
| *Juncus pallidus* | | | | | 1 | 1 | 2012 |
| *Kennedia prostrata* | 1 | | | 1 | | | 2013 |
| *Kunzea glabrescens* | 1 | | | 1 | 1 | | 2015 |
| *Laxmannia ramosa* | 1 | | | | | | 2013 |
| *Laxmannia squarrosa* | 1 | | | | | | 2013 |
| *Lechenaultia floribunda** | 1 | | | 1 | 1 | | 2013 |
| *Lepidosperma* sp. | | | | 1 | | | 2014 |
| *Lepidosperma squamatum** | 1 | | | 1 | | | 2013 |
| *Leptomeria empetriformis* | 1 | | | | | | 2016 |
| *Leucopogon conostephioides* | 1 | | | | | | 2013 |
| *Levenhookia stipitata* | 1 | | | | | | 2012 |
| *Lobelia tenuior* | 1 | | | | 1 | | 2012 |
| *Lomandra caespitosa* | 1 | | | 1 | | | 2014 |
| *Lomandra hermaphrodita* | 1 | | | 1 | | | |
| *Lomandra nigricans* | | | | 1 | | | |
| *Lomandra preissii* | | | | 1 | | | |
| *Lomandra suaveolens* | 1 | | | 1 | | | 2014 |
| *Lyginia barbata/imberbis* | 1 | | | 1 | | | 2015 |
| *Macarthuria apetala* | 1 | | | | | | 2015 |
| *Macarthuria australis* | 1 | | 1 | | 1 | | 2012 |
| *Macrozamia fraseri* | | | 1 | | | | |
| *Melaleuca incana* subsp. *nana* | | | | 1 | | 1 | 2016 |
| *Melaleuca preissiana* | | | | 1 | | 1 | 2015 |
| *Melaleuca rhaphiophylla* | | | | 1 | | 1 | 2017 |
| *Melaleuca seriata* | | | 1 | 1 | | | 2013 |
| *Melaleuca teretifolia* | | | | 1 | | 1 | |
| *Melaleuca thymoides* | 1 | 1 | 1 | 1 | | | 2015 |

| Species | Topsoil | Topsoil and other | Direct Seeded | Planted 2012 -2016 | Local Opportunist | Dampland ecotone only | First started flowering |
|---|---|---|---|---|---|---|---|
| *Melaleuca viminea* | | | | 1 | | 1 | |
| *Microtis media* | | | | | 1 | | 2013 |
| *Millotia tenuifolia* | 1 | | | | | | 2013 |
| *Nuytsia floribunda* | | | 1 | 1 | | | |
| *Orthrosanthus laxus* | | | | 1 | | | |
| *Patersonia occidentalis* | 1 | 1 | 1 | | | | 2013 |
| *Pericalymma ellipticum* | | | | 1 | | 1 | 2017 |
| *Persoonia saccata* | 1 | | | | | | 2016 |
| *Petrophile linearis* | | | 1 | 1 | | | 2015 |
| *Phlebocarya ciliata** | 1 | | | 1 | | | 2016 |
| *Phlebocarya filifolia* | | | | 1 | | | 2017 |
| *Phyllangium paradoxum* | 1 | | | | | | 2012 |
| *Phyllanthus calycinus* | 1 | | | | | | 2013 |
| *Philotheca spicata* | 1 | | | | | | 2016 |
| *Platysace filiformis* | 1 | | | | | | 2013 |
| *Podotheca angustifolia* | | | | | 1 | | 2013 |
| *Podotheca gnaphalioides* | 1 | | | | 1 | | 2012 |
| *Poranthera microphylla* | 1 | | | | | | 2012 |
| *Poranthera moorokatta* | 1 | | | | | | 2012 |
| *Poranthera huegelii* | 1 | | | | | | 2017 |
| *Pultenaea reticulata* | | | 1 | | | 1 | 2016 |
| *Quinetia urvillei* | 1 | | | | | | 2012 |
| *Regelia ciliata* | | | | 1 | | 1 | 2015 |
| *Regelia inops* | | | | 1 | | | 2015 |
| *Rhodanthe citrina* | 1 | | | | | | 2012 |
| *Scaevola repens* | 1 | | | | | | 2015 |
| *Schoenus curvifolius* | 1 | | | 1 | | | 2014 |
| *Schoenus caespititius* | 1 | | | 1 | | | 2014 |
| *Scholtzia involucrata** | 1 | | 1 | 1 | | | 2014 |
| *Senecio condylus* | | | | | 1 | | 2013 |
| *Siloxerus humifusus* | 1 | | | | 1 | | 2012 |
| *Sowerbaea laxiflora* | 1 | | | | | | |
| *Stirlingia latifolia* | 1 | 1 | 1 | 1 | | | 2015 |
| *Stylidium araeophyllum* | 1 | | | | | | 2013 |
| *Stylidium piliferum* | 1 | | | | | | 2013 |
| *Stylidium repens* | 1 | | | | | | 2015 |
| *Synaphea spinulosa* | 1 | | | | | | 2014 |
| *Thysanotus arbuscula* | 1 | | | | | | 2014 |
| *Thysanotus manglesianus* (sp. climbing) | 1 | | | | | | 2015 |
| *Thysanotus arenarius* | 1 | | | | | | 2016 |
| *Thysanotus sparteus* | 1 | | | | | | 2016 |
| *Thysanotus thyrsoideus* | 1 | | | | | | 2013 |
| *Trachymene pilosa* | 1 | | | | | | 2012 |
| *Tricoryne tenella* | 1 | | | | | | |
| *Wahlenbergia preissii* | 1 | | | | | | 2013 |
| *Xanthorrhoea preissii* | | | 1 | 1 | | | |
| *Xanthosia huegelii* | 1 | | | | | | 2013 |
| **TOTAL** | **115** | **15** | **37** | **69** | **22** | **13** | |
