## Supplementary material for "Several complementary methods of plant establishment are required for effective restoration of banksia woodland": Supplimental table S2

**Table S2.** List of open access documents containing additional information about this project. These are available from https://library.dbca.wa.gov.au.


Brundrett M. 2012. *Banksia Woodland Restoration Project Annual Report 2012.* Department of Parks and Wildlife, Crawley, Western Australia. July 2012.

Brundrett M, Clarke K, Longman V. 2012. Setting comprehensive and effective completion criteria for banksia woodland restoration. *Society for Ecological Restoration Australasia Conference, Perth*.

Brundrett M, Clarke K, Longman V. 2014. Setting comprehensive and effective monitoring targets for banksia woodland restoration and management. In: Mucina L, Price JN & Kalwij JM (eds). Biodiversity and Vegetation Patterns, Processes, Conservation. Kwongan Foundation, Perth, Australia, p. 72.

Brundrett M, Clarke K, Taylor K and Wisolith A. 2013. *Banksia Woodland Restoration Project Annual Report 2013.* Department of Parks and Wildlife, Crawley, Western Australia. December 2013.

Brundrett M, Collins, M, Clarke K. Longman V, Wisolith A. 2017. *Flora and Vegetation Completion Criteria*. Department of Biodiversity, Conservation and Attractions, Perth, Western Australia.

Brundrett M, Longman V, Wisolith A, Jackson K and Clarke K. 2018. *Banksia Woodland Restoration Project Annual Report 2017.* Department of Biodiversity, Conservation and Attractions, Perth, Western Australia. 59 pages.

Brundrett M, Longman V, Wisolith A, Jackson K, Collins M and Clarke K. 2016. *Banksia Woodland Restoration Project Annual Report 2016.* Department of Parks and Wildlife, Perth, Western Australia. January 2016.

Brundrett M, Longman V, Wisolith A, Moore T, Taylor K and Clarke K. 2015. *Banksia Woodland Restoration Project Annual Report 2015.* Department of Parks and Wildlife, Crawley, Western Australia. March 2015.

Brundrett M, Longman V, Wisolith A, Taylor K and Clarke K. 2014. *Banksia Woodland Restoration Project Annual Report 2014.* Department of Parks and Wildlife, Crawley, Western Australia. March 2015.

Brundrett M, Van Dongen R, Huntley B, Tay N, Longman, VA. 2019. A monitoring toolkit for banksia woodlands: comparison of different scale methods to measure the recovery of vegetation after fire. *Remote Sensing in Ecology and Conservation* 5(1): 33-54

Brundrett M, Wisolith 2020. *Identification Guide to Seedlings of Banksia Woodland Plants.* Department of Parks and Wildlife, Perth, Western Australia.

Brundrett M, Wisolith A, Collins M, Longman V, Clarke K (2019). Several complementary plant establishment methods are required to restore banksia woodland. In: Commander L (ed). *Nature City Seminar: Book of Abstracts*. South Perth, Australia, p. 6

Clarke, K., Glossop, B., Brundrett, M. and Collins, M. 2017. *Site Selection for Topsoil Transfer and Management Actions*. Department of Biodiversity, Conservation and Attractions, Perth, Western Australia.

Crawford A, Monaghan A, Brundrett M, Wisolith A, Cochrane A (2014). Seed science to improve restoration (ABSTRACT). In 2nd Conference of SERA, Society for Ecological Restoration, From Large to Small Islands, 17-21 Nov 2014, Nouméa, New Caledonia, Nouvata Park: Conference Proceedings p. 84

Keighery G, Longman V, Brundrett M. 2017. Weedy and natural distribution of Acacia trigonophylla (Fabaceae). *Western Australian Naturalist* 31: 53-62.

Longman V, Brundrett M, Wisolith A, Clarke K. 2018. *Banksia Woodland Restoration Project Flora Survey Quadrat Data 2011 – 2017.* Metadata and data file. Department of Biodiversity Conservation and Attractions, Crawley, Western Australia. NatureMap website (url: https://naturemap.dbca.wa.gov.au/).
